# Exploratory multi-omics analysis reveals sex-specific differences in microbial response to antibiotic exposure

**DOI:** 10.64898/2026.09.01.748584

**Authors:** Avani Tantry, Weimiao Long, Julielam Tran, Sidhant Rohatgi, Erika L. Cyphert

## Abstract

Antibiotic exposure is a major driver of microbiome disruption and antimicrobial resistance gene (ARG) expansion. Yet, the role of biological sex in shaping these responses remains poorly understood. Most studies do not stratify antibiotic-induced microbiome changes by sex or integrate multi-omics datasets, limiting our understanding of how microbial, metabolic, and immune responses interact. Therefore, there remains a critical need for an integrative systems-level approach to determine how sex-specific disruptions under antibiotic pressure are paralleled across microbial, metabolic, and host immune layers. The objective of this work was to perform an exploratory study investigating how continuous antibiotic exposure reshaped the gut microbiome across sexual maturation and how these perturbations influenced downstream host responses in a sex-specific manner using an integrative multi-omics framework. Male and female mice that were exposed to continuous antibiotics were profiled over sexual maturation using shotgun metagenomics, untargeted metabolomics, and bulk RNA sequencing of the spleen to assess microbial composition, ARG dynamics, metabolic profiles, and immune responses. Overall, our results demonstrated sex-specific correlations at a systems-level that help provide valuable context to the differences observed in males and females upon antibiotic exposure.

**Importance:** Antimicrobial resistance is a major global threat, and it remains unclear how continuous antibiotic exposure shapes the gut microbiome across sexual maturation and influences the host response in a sex-specific manner. In this work, we used an integrative multi-omics framework to demonstrate sex-specific correlations at a systems-level across several biological layers including taxonomic and resistome composition, host-microbe interactions through cecal metabolomic profiles, and host splenic gene expression. These results provide valuable context to understand systems-level differences in males and females following antibiotic exposure and inform sex-specific antibiotic strategies to contribute to more effective antimicrobial stewardship.

## Introduction

Antimicrobial resistance is a major global threat to human health, with the misuse of antibiotics serving as a key driver in the emergence of drug-resistant pathogens, contributing to approximately 1.5 million deaths annually ^1,2^. Several animal and human studies have shown that biological sex influences antibiotic resistance gene (ARG) burden and microbiome composition across different experimental settings ^3–6^. Moreover, there is growing evidence that biological sex shapes host metabolism, immune response, and host-microbiome interactions.

Antibiotic exposure imposes selective pressure that enables resistant microbes to survive and proliferate, contributing to the emergence of antimicrobial resistance ^2,7^. Collectively, these resistance genes that are present in microbial communities are termed the “resistome” ^8^. In addition to altering resistance dynamics, antibiotic-induced changes to the gut microbiota also contribute to substantial changes in metabolic processes and corresponding metabolites produced by the host ^9,10^. This alteration of metabolic activity may also be associated with host inflammatory responses in the colon such as up-regulation in gene expression of various pro-inflammatory cytokines such as IFN-γ, IL-1β, and IL-6 ^10^. While many common infections have an antibiotic course lasting less than 14 days, prolonged antibiotic therapy lasting several weeks may be required for chronic or severe infections, prophylaxis in immunocompromised patients, or as adjunctive therapy in inflammatory conditions ^11,12^. Although short-term antibiotic exposure is associated with a profound loss of diversity and shift in community composition, partial recovery toward baseline can occur within weeks following antibiotic cessation, although this recovery is often incomplete ^13^. Despite the clinical use of prolonged antibiotic therapy, the effects of sustained antibiotic exposure on the gut microbiome and downstream host outcomes remain comparatively underexplored ^12,14^.

Research across human and animal studies has demonstrated that biological sex plays a crucial role in shaping the composition and diversity of the gut microbiota, host metabolism, and immune responses ^15–18^. Though biological sex is known to affect gut microbiota composition, relatively few studies have investigated whether antibiotics elicit sex-specific differences in the gut microbiome and how it alters microbial and host metabolism, and immune responses ^19–21^. While age related sex differences in the gut microbiota are well studied in early childhood and adulthood, comparatively little is known about these differences during puberty, a period of considerable developmental and hormonal changes ^16^. Emerging evidence suggests that sex-specific microbiota trajectories may become more pronounced during this developmental window, which may be particularly relevant given that antibiotic exposure during pubertal age can induce long lasting alterations to the gut microbiota composition ^22–24^. Despite prior evidence suggesting sexually dimorphic responses to antibiotic exposure, a crucial gap is that many studies that collectively investigated microbial, metabolic, or host immune consequences of antibiotic perturbation used single-sex cohorts or analyzed aggregated data without sex stratification potentially masking biologically meaningful sex-specific outcomes ^10,25–28^. There is a critical need for an integrative systems-level approach to determine how sex-specific disruptions under antibiotic pressure are paralleled across microbial, metabolic, and host immune layers. This knowledge gap is particularly relevant during developmental stages such as puberty, when both the gut microbiome and host immune system undergo substantial maturation. Understanding these sex differences is therefore valuable to enable sex-aware microbial therapeutics and to inform sex-specific antibiotic strategies resulting in more effective antimicrobial stewardship ^19^.

The objective of this work was to perform an exploratory study investigating how continuous antibiotic exposure reshaped the gut microbiome across sexual maturation and how these perturbations influenced downstream host responses in a sex-specific manner using an integrative multi-omics framework. C57Bl/6 mice were used, that reach sexual maturity from 6-8 weeks of age ^29^. The study was designed to capture antibiotic perturbations across this developmental transition, with sample collections structured around a four-week baseline timepoint (before antibiotic exposure), a five-week pre-sexual maturity stage, and a 10-week post-sexual maturity stage. Specifically, this work aimed to 1) evaluate how biological sex shaped antibiotic-driven remodeling of the gut microbiome across sexual maturation and 2) evaluate how antibiotic-associated shifts in the gut microbiome, resistome, and metabolome related to sex-dependent splenic immune transcriptional responses using integrated multi-omic analyses.

## Materials and Methods

### Mouse strains

All animal procedures received prior approval from the local Institutional Animal Care and Use Committee (IACUC). Male and female C57Bl/6 mice were purchased from Jackson Laboratory (Sacramento, CA, USA) and bred using trio breeding. Pups obtained from the first breeding cycle were used for this experiment. Pups were weaned at three weeks of age where they were separated by sex and redistributed across cages (n=5/cage), to minimize litter-specific and cage-specific environmental effects. All cages contained corn cob bedding (Envigo Sani-Chip 7090A standard, CA, USA), standard laboratory chow (8604 Rodent Diet, Harlan Teklad, USA), deionized (DI) water *ad libitum* which was filtered and treated with bleach for sterilization, as well as cotton based nestlets for enrichment (Envigo Nestlets, CA, USA).

### Experimental design and antibiotic dosing

A primary cohort of male and female pups were each divided into Control and Antibiotic groups: F_control, F_abx, M_control, M_abx (n=5/group/sex) (**Figure 1A**). To enable endpoint sample collection at the earlier developmental stage without disrupting the primary cohort, a secondary cohort with an identical experimental design was established with an endpoint of five weeks of age. Mice in the Antibiotic group were dosed with an antibiotic cocktail consisting of 0.5 g/L neomycin (Sigma Aldrich, USA, Catalog #: 102706992) and 1 g/L ampicillin (Sigma Aldrich, USA, Catalog #: 0000442939) administered via their drinking water, whereas the Control groups received standard drinking water. To prevent light-induced degradation of antibiotics, water bottles for the Antibiotic group were wrapped in aluminum foil and further covered with polypropylene material. Antibiotic dosing was initiated at four weeks of age and lasted for the duration of the experiment until the endpoint of the primary cohort at 10 weeks of age. The antibiotic cocktail was replaced every 2-3 days over the duration of the study. Ampicillin and neomycin were selected due to their poor oral bioavailability and capability of acting directly on gut microbes, while minimizing systemic exposure. Three biological replicates per group were randomly selected for inclusion in the study analyses.

**Figure 1.**
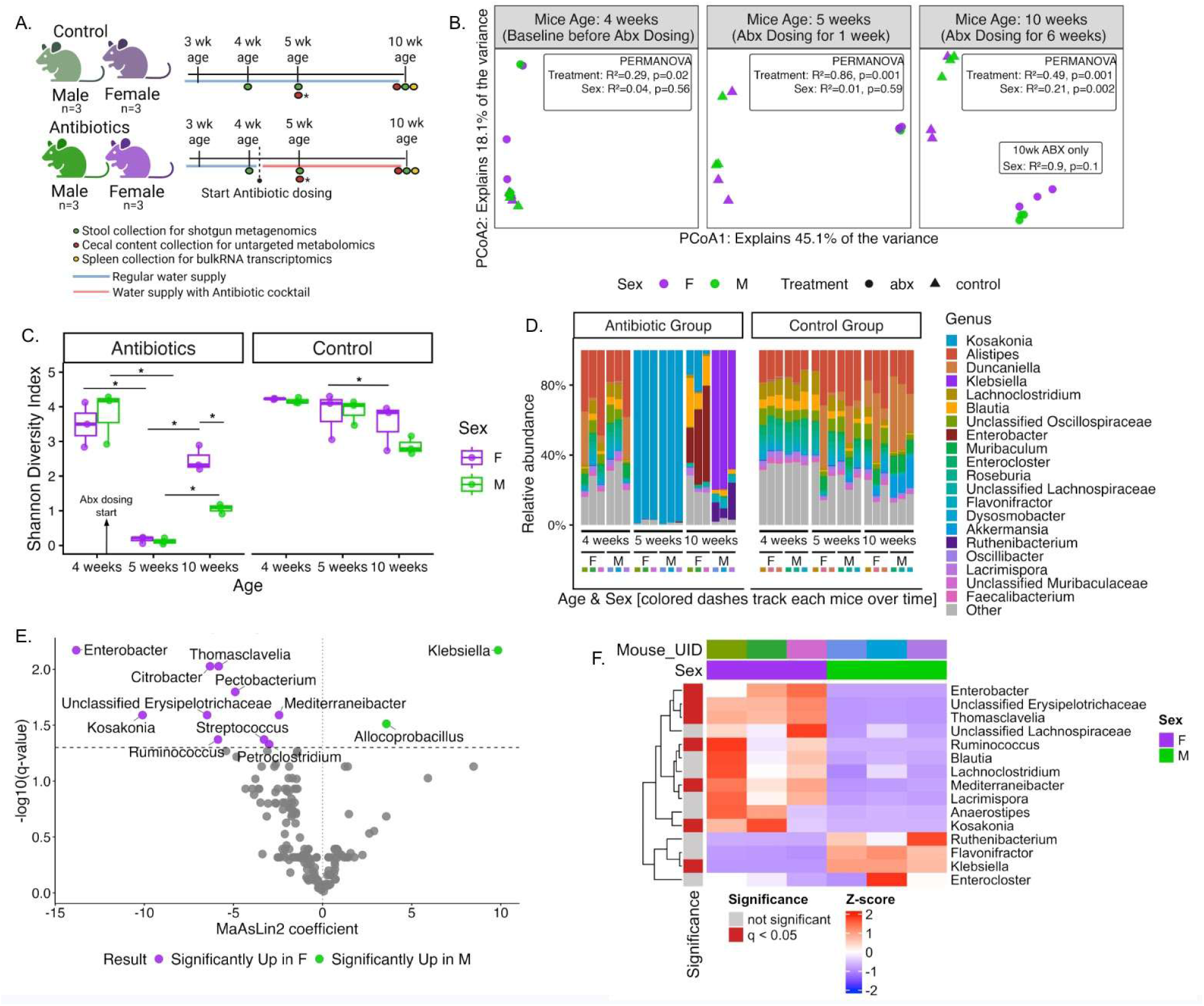
**(A)** Experimental workflow of study. **(B)** Bray-Curtis PCoA of taxonomic composition across samples stratified by age, sex, and treatment revealed divergence in taxonomic structure between F_abx and M_abx at 10 weeks of age. **(C)** Shannon diversity of microbial taxonomy across age, sex, and treatment groups revealed that females gained a significantly higher diversity than males upon antibiotic exposure at 10 weeks of age (* = BH-adjusted p-value < 0.05). **(D)** Relative abundance of top 20 most abundant genera over time differed by sex in antibiotic exposed mice at 10 weeks of age. **(E)** MaAsLin2 volcano plot of genera between M_abx and F_abx at 10 weeks of age demonstrated distinct sex-specific microbial community structure following six weeks of antibiotic exposure. **(F)** MaAsLin2 heatmap of top 15 most abundant genera in M_abx and F_abx at 10 weeks of age. Genera with a mean relative abundance < 1% were excluded from the analysis.

### Stool sample collection

Fresh stool samples were collected from all groups in the primary cohort at four weeks of age (baseline before antibiotic exposure), five weeks of age (one week of antibiotic exposure), and 10 weeks of age (endpoint, six weeks of antibiotic exposure). Mice were placed in sterile plastic containers and stool samples were collected in sterile cryotubes (CellPro VC118-C 2.0 mL cryovials, Alkali Scientific, USA). Stool samples were transported to −80°C freezer for long-term storage. Stool samples were consistently collected by 8:00 AM – 11:00 AM at all timepoints to minimize the effects of circadian rhythm on microbiota composition ^30^. Mice weight was also recorded at the time of stool sample collection and monitored across the duration of the study (**Supplemental Figure 1**).

### Spleen and cecal content collection

Upon reaching the experimental endpoints for the secondary cohort (5 weeks of age) and primary cohort (10 weeks of age) the spleen and cecal contents were collected. The spleens were harvested by positioning mice in a right lateral recumbency to expose the left side and performing a vertical side incision on the left flank directly over the spleen to expose the organ for excision. Following excision, spleens were immediately submerged into a pre-aliquoted RNAlater solution (Sigma Aldrich, USA) in sterile cryosafe tubes and immediately placed on ice. Spleen samples were temporarily stored in the 4°C refrigerator until ready for bulk RNA sequencing. Cecal contents were harvested by exposing the cecum via a mid-abdominal incision and carefully puncturing the end to provide an opening. Cecal contents (chyme) were extracted directly into sterile cryosafe tubes without the inclusion of cecal wall tissue to isolate the luminal metabolites. The tubes were immediately placed on dry ice and later stored in a −80°C freezer until ready for untargeted metabolomics.

### Shotgun metagenomics library preparation and sequencing

Shotgun metagenomic DNA extraction, library preparation, and sequencing were performed by the UC San Diego Microbiome Core following previously published protocols^31^. Briefly, nucleic acid extraction and purification from samples was conducted using the MagMAX Microbiome Ultra Nucleic Acid Isolation Kit (Thermo Fisher Scientific, USA) and automated on the KingFisher Flex robotic platform (Thermo Fisher Scientific, USA). Blank controls (negative controls) and mock communities (positive controls) (Zymo Research Corporation, USA) were included on each extraction plate and were carried through all downstream processing steps. DNA concentration was quantified using a PicoGreen fluorescence assay (Thermo Fisher Scientific, USA), and metagenomic libraries were prepared using KAPA HyperPlus kits (Roche Diagnostics, USA) in a miniaturized one-fifth reaction volume format and automated on the EpMotion automated liquid handlers (Eppendorf, Germany). Sequencing was performed at the Institute for Genomic Medicine (IGM), UC San Diego on an Illumina NovaSeq X Plus platform with paired-end 150 bp cycles and a sequencing depth of 5 million reads.

### Untargeted metabolomics liquid chromatography-mass spectrometry (LC-MS)

Liquid chromatography mass spectrometry procedures were outsourced to the UC San Diego Microbiome Core. Cecal samples were removed from −80°C, thawed, and ∼20 mg sample was placed into round-bottom Eppendorf tubes (Catalog #: 0030123620). A single steel bead (Qiagen Stainless Steel Beads, 5 mm Catalog # 69989) was added to the sample tube and 800 μL of ice-cold extraction solvent was added to it, maintaining a 1:40 sample to solvent ratio (extraction solvent – 50:50 LC/MS grade methanol Fisher Scientific Catalog # A456-4, LC/MS grade water Fisher Scientific Catalog # W6-4). The samples were then homogenized at 25 Hz for five minutes using a Qiagen Tissuelyzer and then incubated at 4°C for 30-60 minutes until precipitation. Once precipitated, the samples were centrifuged at > 15000 g for 10 minutes. The supernatant was carefully transferred to a new 96 well-plate and was vacuum concentrated to dryness via centrifugal lyophilization (Labconco Centrivap, USA). The dried samples were then stored at −80°C until LC-MS was performed.

For the LC-MS run, the dried samples were resuspended with 200 μL of methanol:water (1:1) containing 1 μM sulfadimethoxine as an internal standard. Untargeted metabolomics was performed using a Vanquish liquid chromatography system (Thermofisher Scientific, USA) coupled to a QExactive mass spectrometer (Thermofisher Scientific, USA) with a C18 column (Phenomenex Kinetex 1.7 μm C18 100 Å LC Column 50 x 2.1 mm). The mobile phase used was LC-MS grade water (phase A) and LC-MS grade acetonitrile (phase B), both containing 0.1% formic acid (Fisher Scientific, Optima LC-MS), with a flow rate set to 0.5 mL/min. Samples were injected at 95% A: 5% B, and were held for one minute, before ramping up to 100% B over seven minutes, which was then held for 0.5 minutes before returning to starting conditions. Data was collected in positive ion mode using data-dependent acquisition to acquire MS full scan spectra, followed by MS/MS spectra of the top five most abundant ions.

### Bulk RNA transcriptomics library preparation and sequencing

RNA extraction and library preparation were performed by the Sanford Consortium for Regenerative Medicine (SCRM) Genomics Core. Total RNA was extracted using the Qiagen RNeasy Plus Universal Kit (Catalog # 73404) following homogenization with the Qiagen TissueRuptor II according to manufacturer protocol. RNA concentration was quantified using the ThermoFisher Qubit 3.0 Fluorometer with the Qubit RNA Broad Range Assay Kit (Catalog # Q10211), while RNA purity was assessed using the ThermoFisher NanoDrop 2000 Spectrophotometers. RNA integrity, determined via an RNA integrity number (RIN), was evaluated on the Aligent TapeStation 4200 using the RNA ScreenTape Assay. The RIN was determined to be above 9 for all samples. Following the initial quality control assessment, 1000 ng of total RNA was converted to an Illumina sequencing library, with library preparation performed using the Illumina Stranded mRNA Prep kit (Catalog # 20040534) according to manufacturer protocol. The resulting libraries were quantified on the ThermoFisher Qubit 3.0 Fluorometer using the 1x dsDNA High Sensitivity Assay Kit (Catalog # Q33231). Library size was determined on the Aligent TapeStation 4200 using the D1000 ScreenTape Assay. Libraries were normalized to 10 nM, pooled at equal volumes, and subsequently sequenced on a single lane of an Illumina NovaSeq X Plus 10B flow cell using a paired-end 100 bp (PE100) sequencing configuration with an expected sequencing depth of about 100M reads per sample.

### Shotgun metagenomics data preprocessing

The paired end .fastq files generated from shotgun metagenomics sequencing were preprocessed using the open-source Chan Zuckerberg ID (CZID) platform ^32^. Quality control preprocessing within CZID included sequence quality filtering using Fastp, host read subtraction using Bowtie2 and Hisat2 alignment against the mouse reference genome, duplicate read removal, and random subsampling to 1 million fragments prior to downstream taxonomic classification and antimicrobial resistance gene identification (**Supplemental Table 1**). For taxonomic classification mNGS pipeline version 8.3 was used, and for antimicrobial resistance gene identification AMR pipeline version 1.4.2, CARD Database version 3.2.6, and Wildcard Database version 4.0.0 was used. Microbial reads were aligned to the NCBI NT database (National Center for Biotechnology Information Nucleotide Database) using minimap2 (assembly-based alignment). Microbial reads were also aligned to the NCBI NR database (non-redundant protein) using Diamond.

Sample taxon reports were exported from CZID. For both genus and species level bacterial annotations, taxonomic read assignments were filtered per CZID’s recommended criteria to retain a nucleotide reads per million (nt_rpm) greater than 10, nucleotide alignment length of greater than 50 base pairs, and non-zero nucleotide (nt) and normalized (nr) read counts. Following filtration species/genus names and corresponding nt_rpm values were normalized using the total sum scaling (TSS) method to obtain relative abundances used for downstream analyses. In parallel, for antimicrobial resistance analysis, combined AMR reports were exported from CZID. ARG annotations were filtered to only retain features with a read coverage breadth greater than 5% prior to downstream analyses.

Microbial functional pathway abundance profiles were generated using the bioBakery WGS pipeline (User options: UniRef level = UniRef90; UniRef group = KEGG orthology (KO); units = copies per million, run StrainPhlAn = not chosen) and was implemented in Nephele (National Institute of Allergy and Infectious Diseases (NIAID) Office of Cyber Infrastructure and Computational Biology (OCICB) in Bethesda, MD) ^33^. Paired-end FASTQ files generated by CZID pipeline following its mouse host filtering step were used as input since the Nephele bioBakery pipeline does not include a mouse reference for host read removal.

### Untargeted metabolomics data preprocessing

Raw QExactive files obtained from the LC-MS run were converted to mzML format using the MSConvert software in the ProteoWizard tool. These reformatted files were then imported and processed in MZmine (v4.7.8) for filtering and feature finding. Mass detection was performed on both MS1 and MS2 spectra using Orbitrap-appropriate intensity thresholds and chromatogram construction was performed using ADAP (Automated Data Analysis Pipeline) chromatogram builder. Features were detected using the ADAP local minimum deconvolution algorithm, isotopic peaks were grouped and filtered to remove redundancy, and features were aligned across samples using the join aligner. Post-alignment processing included row filtering and duplicate peak removal. Finally, correlation-based grouping (metaCorrelate), ion identity networking, and lipid annotations were applied. The resulting feature list was exported as a feature quantification table (.csv) and MS/MS spectral summary (.mgf) for downstream analysis and submission to Global Natural Products Social Molecular Networking (GNPS) for feature based molecular networking.

The feature quantification table (.csv) and MS/MS spectral summary (.mgf) generated from MZmine were submitted to GNPS2 for feature-based molecular networking using the standard GNPS2 workflow. Spectral data were filtered and matched against public spectral libraries using a fragment ion tolerance of 0.02 Da, with library matches retained at a minimum cosine similarity score of 0.7 and at least five matched fragment peaks. Molecular networks were constructed based on pairwise spectral similarity using the same cosine threshold (0.7) and minimum matched peaks (n=5), with a maximum connected component size of 100 nodes and top 10 edges retained per node to control network complexity. No additional normalization or precursor filtering was applied within GNPS. Library searching incorporated multiple GNPS spectral collections, including propagated and curated metabolomics libraries. The resulting molecular networks, spectral annotations, and cluster summaries were generated through the GNPS2 pipeline and used for downstream metabolite annotation and interpretation.

### Bulk RNAseq transcriptomics data preprocessing

Bulk RNAseq data was preprocessed using a standardized pipeline which included genome indexing, read quality control, alignment, and gene-level quantification. A reference genome sequence (FASTA) and its corresponding gene annotation file (GTF) for *Mus musculus* was obtained from Ensembl (GRCm39 build) and used to generate a genome index. Both files were derived from the same version to maintain compatibility. A genome index was built using the tool STAR (Spliced Transcripts Alignment to a Reference) ^34^. The obtained raw paired-end FASTQ files were assessed for sequencing quality metrics using FastQC and were subsequently trimmed using fastp (v0.23.2) to remove low-quality bases and adapters, generating cleaned paired-end FASTQ files. The trimmed reads were then aligned to the built genome index using STAR in two-pass mode to improve splice junction detection. Alignments were saved as coordinate-sorted Binary Alignment Map (BAM) files and gene-level read counts were computed for all samples from their BAM files using featureC ounts which included exon-level counting and gene_id annotation (**Supplemental Table 2**). The resulting tab-separated file (.tsv) was converted to comma-separated (.csv) format for compatibility with downstream analyses. Finally, genes with zero counts across all samples were removed, and additional prefiltering was performed where only genes with at least 10 counts in a minimum of three samples were retained, resulting in a filtered dataset of approximately 28,000 genes for downstream analysis.

### Shotgun metagenomics data analysis

To evaluate changes in microbiota composition longitudinally, alpha and beta diversity were calculated across all samples using the vegan package in R (v 2.7.3) ^35^. Specifically, Shannon and Simpson diversity metrics were calculated. Welch’s t-tests were performed to evaluate significance in sex differences, and paired t-tests were used to evaluate significance across ages. Bray-Curtis principal coordinate analysis was performed followed by PERMANOVA. The relative abundance of the top 20 most abundant genera across all samples were evaluated to assess community composition. To further evaluate the effect of age and sex on an antibiotic disrupted gut, univariate analysis was performed using the MaAsLin2 package in R (v 4.0.2) ^36^. The TSS normalized data was log transformed for the MaAsLin2 analysis and the resulting data was visualized using volcano plots and heatmaps.

ARG abundance was compared across samples by summing the dpm (depth per million) values of all detected ARGs post-filtration in each sample. ARG richness across samples was compared by summing the number of unique ARGs with non-zero dpm values in each sample. Welch’s t-tests were performed to evaluate the significance in sex differences and paired t-tests were used to evaluate significance across ages. The data was then TSS normalized and Bray-Curtis principal coordinate analysis was performed followed by PERMANOVA to compare ARG composition across samples. Resistance genes were then categorized per their CARD (Comprehensive Antibiotic Resistance Database) high level drug class annotations and stacked relative abundance plots were used to visualize resistance gene composition at drug class level. To further evaluate the effect of age and sex on an antibiotic disrupted resistome, univariate analysis was performed using the MaAsLin2 package in R ^36^. The TSS normalized data was log transformed for the MaAsLin2 analysis, and the resulting data was visualized using volcano plots and heatmaps.

Microbial functional pathway-level analyses used HUMAnN pathway abundances (copies per million) converted to relative abundance. To evaluate the effect of age and sex on microbial pathways in an antibiotic disrupted microbiome, univariate analysis was performed using the MaAsLin2 package in R ^36^. The TSS normalized data was log transformed for the MaAsLin2 analysis, and the resulting data was visualized using volcano plots and heatmaps.

### Untargeted metabolomics data analysis

The final metabolomics feature table and metabolite annotations were obtained from the GNPS platform following feature based molecular networking analysis. Raw metabolomic features were subsequently subjected to a series of quality control and filtration steps. Features with a pool-to-blank ratio less than five were removed to minimize background contamination. Retention time filtering was then applied to exclude features with retention times below 0.2 minutes or above eight minutes. Then, features containing zero intensity values across all biological samples were removed, followed by prevalence filtering in which only features present in at least two-thirds of samples within at least one experimental group were kept. Finally, a sixmix quality-control filter was applied, where features with a pool-to-sixmix ratio of less than five were removed.

The features that remained post QC in the biological samples were TIC (Total Ion Current) normalized and rCLR (robust Centered Log Ratio) transformed followed by minimum value imputation for the zero values^37^. The TIC-rCLR normalized data was then used for downstream analysis. Principal component analysis (PCA) followed by PERMANOVA was performed to analyze sample level differences in metabolomic composition. UpSet plots were also used to look for shared and unique metabolite composition. To identify features driving separation between groups, Partial Least Squares Discriminant Analysis (PLS-DA) followed by a linear regression was performed. From the PLS-DA analysis, only features with a Variable Importance in Projection (VIP) score greater than one and an absolute Cliff’s delta value of greater than 0.5 were selected for the linear regression analysis. From the linear regression results only annotated features with a Benjamini-Hochberg (BH) adjusted p-value of 0.05 were considered for further analysis. PLS-DA was performed on the 10 weeks of age samples in the Antibiotic groups and results from this analysis were used to perform a linear regression using sex as the variate. Selected features from this analysis were visualized using heatmaps and volcano plots. Putative annotations of selected features were manually cleaned (**Supplemental Table 3**).

### Bulk RNAseq transcriptomics data analysis

To identify differentially expressed genes (DEGs) from the bulk RNAseq data, DESEQ2 was performed and volcano plots were used to visualize the data. To evaluate how males and females responded differentially to antibiotics relative to their respective controls, DESEQ2 was first performed using ∼Treatment*Sex as the model design. To further evaluate sex-specific differences between males and females in response to antibiotics, DESEQ2 was also performed on just the antibiotic group using ∼Sex as the model design. For functional analysis, only the DEGs that had a |log_2_ fold change| > 1 and an adjusted p-value of < 0.05 were retained. The clusterProfiler package in R (v 4.14.6) was used to perform functional enrichment analysis, and enrichment results were represented using dotplots.

### Spearman correlation analysis

To identify correlations between specific features within any two omics datasets, Spearman correlation analysis was performed. Specifically, correlation was assessed between taxa and ARGs, taxa and functional pathways within shotgun metagenomics data. Correlations were also assessed between microbes and metabolites and functional pathways and metabolites in a metagenomics-metabolomics Spearman correlation analysis. The corrplot() package in R (v 0.95) was used to visualize the Spearman correlation results.

### Multi-omics integration analysis

The QC filtered, normalized, and processed taxa, ARG, metagenomics functional pathways, metabolomics, and RNAseq matrices were integrated for a multi-omics analysis using the DIABLO (Data Integration Analysis for Biomarker discovery using Latent cOmponents) tool within the mixOmics package in R (v 6.30.0) ^38^. A DIABLO design = 0.5 was chosen for this analysis to strike a balance between correlation and discrimination between datasets. The model was optimized to perform an N-integrative sparse Partial Least Squares Discriminant Analysis (sPLS-DA) using the block.splsda tool. sPLS-DA was performed along two components, and 10 features per component per omic block were retained for the analysis, aiming for high predictive performance with minimal signatures. The results from the DIABLO analysis were then visualized using block PLS-DA plots and corresponding loading plots for each omics block, followed by a clustered image map (CIM) to assess correlation patterns across omics blocks. CIM performs hierarchical clustering using Euclidean-distance-based similarity structure, so to facilitate interpretation of the CIM structure, Ward hierarchical clustering using Euclidean distance was additionally applied to the DIABLO-selected feature matrix. The results were partitioned into four feature clusters and the dendextend package in R (v 1.19.1) was used to construct dendrograms of the four clusters for downstream visualization and biological interpretation ^39^.

## Results

### Sex-specific trends observed across sexual maturity in taxonomy and ARG composition upon antibiotic exposure

To assess whether antibiotic exposure induced sex-specific shifts in microbial community structure across sexual maturation, a beta diversity analysis was performed using Bray-Curtis dissimilarity and visualized using PCoA. At baseline (four weeks of age), female and male samples showed no specific clustering by sex (PERMANOVA: p = 0.26; M vs F at four weeks of age averaged across treatment groups) (**Figure 1B**). Baseline samples showed separation by treatment (PERMANOVA: p = 0.016, Control vs Antibiotics at four weeks of age averaged across sex). Following one week of antibiotic exposure and before sexual maturity (five weeks of age), F_abx and M_abx significantly separated from baseline and their respective controls (PERMANOVA: p = 0.003, Control vs Antibiotics at five weeks of age averaged across sex) consistent with an antibiotic-induced community disruption. No sex-specific separation was observed. Upon six weeks of antibiotic exposure across sexual maturity (10 weeks of age), a separation between F_abx_ and M_abx groups was observed. While this separation was not statistically significant (PERMANOVA: p = 0.10, F_abx vs M_abx at 10 weeks of age), the divergence suggested sex-specific remodeling of antibiotic-perturbed microbial communities over time across sexual maturation.

To determine antibiotic-associated changes in sample-specific diversity, alpha diversity was assessed using Shannon and Simpson diversity indices. At baseline, the Shannon diversity was comparable across all groups (**Figure 1C**). At five weeks of age, the antibiotic exposure resulted in a drastic reduction in Shannon diversity in both sexes, consistent with an extensive depletion of taxonomic richness following acute antibiotic treatment. At 10 weeks of age, a partial recovery of diversity was observed in both F_abs and M_abx. However, F_abx showed a significantly greater Shannon diversity than M_abx (Welch’s t-test p < 0.05), suggesting a more extensive diversification in F_abx following continuous antibiotic exposure across sexual maturation. Similar trends were observed in the Simpson diversity index (**Supplemental Figure 2**).

Relative abundance profiling of the top 20 most abundant genera shed light into the taxonomic composition underlying the observed diversity shifts (**Figure 1D**). Prior to reaching sexual maturity at five weeks of age, both F_abx and M_abx samples showed almost complete domination by *Kosakonia*, showing evidence of a community and diversity collapse in both sexes. The taxonomic composition diverged in a sex-specific manner after sexual maturation at 10 weeks of age. Specifically, F_abx mice presented with a broader distribution of genera including *Enterobacter*, *Blautia*, *Lachnoclostridium*, *Mediterraneibacter*, while also retaining *Kosakonia*. Conversely, M_abx mice were dominated by fewer genera such as *Klebsiella*, *Ruthenibacterium*, and *Flavonifractor*, almost entirely eliminating *Kosakonia*. Control samples on the other hand maintained a relatively stable and diverse composition across all timepoints, with no dominance by opportunistic taxa.

Statistical validation of the observed sex-differences in taxonomic composition at 10 weeks of age between M_abx and F_abx was calculated using MaAsLin2 ^36^. The volcano plot revealed several genera were significantly enriched in F_abx such as *Kosakonia*, *Enterobacter*, *Thomasclavelia*, *Mediterraneibacter*, and *unclassified_Erysipelotrichaceae*. In contrast, M_abx were significantly enriched with *Klebsiella* and *Allocoprobacillus* (**Figure 1E**). A heatmap visualization of the top 15 most abundant genera revealed that most MaAsLin2 significant taxa were also highly abundant (**Figure 1F**). It also demonstrated that additional genera that were not statistically significant such as *Blautia*, *Lachnoclostridium*, *Lacrimispora*, and *Ruminococcus*, had higher abundance favoring F_abx, and genera such as *Ruthenibacterium*, *Flavonifractor*, and *Enterocloster* showed higher abundance in M_abx. Additional MaAsLin2 analyses evaluating the effect of antibiotics across age in M_abx and F_abx are shown in supplemental figures (**Supplemental Figures 3-6**). In summary, the taxonomic analysis demonstrated that upon continuous antibiotic exposure across sexual maturation, a sex-specific microbial remodeling trajectory was observed after sexual maturation. This occurred following a similar community collapse during early antibiotic exposure prior to sexual maturation. Females showed a greater taxonomic diversification relative to males who were dominated by fewer opportunistic taxa.

In addition to the pronounced antibiotic-associated remodeling observed in microbial taxonomic composition, resistome profiles were analyzed to determine whether antibiotic exposure induced sex-specific differences in ARG burden and composition across sexual maturation. The total ARG burden remained low and relatively stable across all Control groups and in the baseline F_abx and M_abx groups (**Figure 2A**). Following antibiotic exposure, ARG abundance significantly increased in both F_abx and M_abx at five weeks of age relative to baseline (paired t-test p < 0.05), indicating rapid ARG expansion under antibiotic pressure. Continued antibiotic exposure through 10 weeks of age maintained the elevated ARG load in both sexes, however a significant sex-difference was observed in which M_abx exhibited a greater total ARG burden than F_abx mice (p < 0.05). ARG richness data also demonstrated similar trends (**Supplemental Figure 7**).

**Figure 2.**
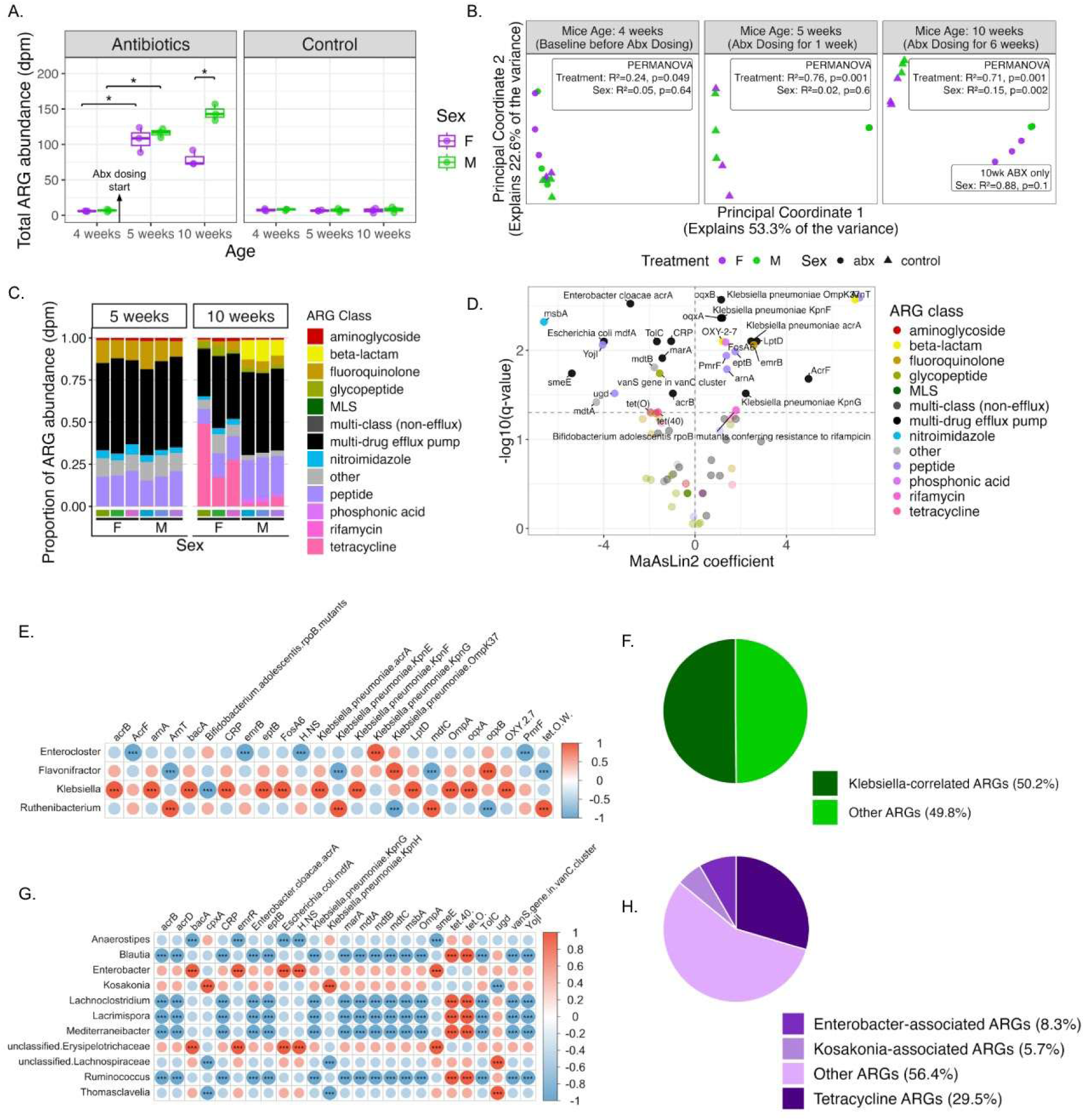
**(A)** Total ARG burden across all samples revealed that M_abx mice had a significantly higher ARG burden following antibiotic exposure at 10 weeks of age relative to F_abx (* = BH adjusted p-value < 0.05). **(B)** Bray-Curtis PCoA of ARG profile revealed divergence of ARG composition between M_abx and F_abx mice at 10 weeks of age. **(C)** Relative abundance of ARGs grouped by high-level drug classes in M_abx and F_abx at 5 and 10 weeks of age. **(D)** MaAsLin2 analysis of ARGs in M_abx and F_abx at 10 weeks of age demonstrated sex-specific ARG structure following six weeks of antibiotic exposure (enriched in females top left, enriched in males top right). **(E)** Spearman correlation analysis between differentially abundant taxa and ARGs in M_abx at 10 weeks of age revealed that ARG composition was strongly associated with *Klebsiella* (*** = BH-adjusted p-value < 0.001). **(F)** Fraction of total ARG abundance represented by genes significantly positively correlated with *Klebsiella* accounted for > 50% of total ARG burden in M_abx at 10 weeks of age. **(G)** Spearman correlation analysis between differentially abundant taxa and ARGs in F_abx at 10 weeks of age revealed that ARG composition was strongly associated with a more diverse range of microbes than M_abx. **(H)** Fraction of total ARG abundance represented by genes significantly associated with dominant taxa in F_abx at 10 weeks of age.

To assess the overall resistome restructuring, Bray-Curtis PCoA was performed on ARG profiles (**Figure 2B**). Prior to antibiotic exposure and in Control samples, resistome profiles demonstrated minimal separation by sex. At five weeks of age (following one week of antibiotic exposure), the antibiotic perturbation became the primary source of variation in ARG compositional structure with limited separation by sex, indicating broadly similar resistome expansion to antibiotic exposure prior to sexual maturation. At 10 weeks of age, a clear divergence between F_abx and M_abx resistomes was observed. However, a PERMANOVA analysis revealed that this observed separation did not reach statistical significance (R^2^ = 0.88, p = 0.1). Together, these findings suggested that sex-dependent resistome remodeling became more apparent after sexual maturation following prolonged antibiotic exposure.

At a high-level drug class, additional insight was gained about the nature of divergence in ARG composition. Prior to antibiotic exposure, ARG profiles in both sexes were dominated by low-abundance tetracycline ARGs (**Supplemental Figures 8-9**). At five weeks of age (following one week of antibiotic exposure), both F_abx and M_abx resistome were dominated by multidrug efflux pump-associated genes, along with peptide and fluoroquinolone resistance genes, demonstrating a similar early adaptation to antibiotic exposure (**Figure 2C**). By 10 weeks of age, sex-dependent shifts in ARG composition were observed. M_abx presented with an increase in beta-lactam specific resistance genes, whereas in F_abx, a higher re-emergence of tetracycline associated resistance genes was observed. These findings suggested that continuous antibiotic exposure was not only associated with an expansion of ARG load, but also with sex-specific restructuring of ARG composition.

This underlying sex-specific compositional divergence in the antibiotic groups at 10 weeks of age was statistically validated to characterize gene-level differences using MaAsLin2 (**Figure 2D** and **Supplemental Figure 10**). M_abx mice showed a significant enrichment of several ARGs associated with beta-lactam resistance, peptide resistance, and multi-drug efflux pumps including *OXY-2-7*, *ArnT*, *ArnA*, *PmrF*, *eptB*, *oqxB*, *Klebsiella pneumoniae KpnF* and *KpnG* and *AcrF*. In contrast, F_abx mice showed enrichment of different multi-drug efflux-associated genes including *smeE*, *Escherichia coli mdfA*, *Enterobacter cloacae acrA*, *TolC*, and *acrB*, nitroimidazole-associated *msbA*, and peptide resistance genes such as *YojI* and *ugd*. No significant sex-associated differences were observed at 5 weeks of age in M_abx and F_abx apart from *acrB* being higher in males, indicating that sex-specific resistome differences emerged after sexual maturation following prolonged antibiotic exposure (**Supplemental Figures 11-12**). Additional MaAsLin2 analyses evaluating the effect of antibiotics across age in M_abx and F_abx are shown in supplemental figures (**Supplemental Figures 13-16**).

Collectively, these findings demonstrated that while antibiotic exposure rapidly expanded ARG burden in both sexes, early resistome responses were broadly similar between sexes. Further, with prolonged antibiotic exposure across sexual maturation, the resistome composition diverged in a sex-associated manner.

To further investigate potential associations between taxa and ARGs, a Spearman correlation analysis was performed among F_abx and M_abx separately at 10 weeks of age using abundant taxa (relative abundance cut-off threshold > 1%) and ARGs that were either significant or abundant (ARGs included were either p < 0.05 MaAsLin2 or among the top 20 most abundant ARGS) from each group. In M_abx, the ARG-taxa correlation revealed *Klebsiella* exhibited significant positive correlations with numerous ARGs, including multidrug efflux pumps *oqxA* and *AcrF*, beta-lactam resistance gene *OXY-2-7*, peptide resistance genes *ArnA* and *eptB*, *Klebsiella* associated genes *KpnF* and *AcrA*, and phosphonic acid resistance gene *FOSA6*. Additional significant positive correlations were also observed with *Flavonifractor* and *Enterocloster* (**Figure 2E**). ARGs that were significantly positively correlated with *Klebsiella* accounted for approximately 50.2% of the total ARG burden in M_abx samples, indicating that a considerable fraction of the male resistome was associated with *Klebsiella* expansion (**Figure 2F**).

Conversely in F_abx, there was a more distributed ARG-taxa correlation structure. Significant positive correlations were observed with tetracycline resistance genes *tet(40)* and *tet(O)* across multiple taxa, including, *Mediterraneibacter*, *Ruminococcus*, *Lachnoclostridium*, *Blautia*, and *Lacrimispora* (**Figure 2G**). These two tetracycline resistance genes made up 29.5% of the total ARG load in F_abx, whereas the resistance genes that had a significant positive correlation with *Enterobacter* and *Kosakonia* were only 8.3% and 5.7% respectively (**Figure 2H**). Collectively, these findings suggested that the male resistome was concentrated around a dominant taxa, whereas the female resistome was suggestive of a broader community level distribution.

### Sex-dependent effects on metabolic remodeling in response to antibiotic exposure

The functional potential of the microbes present in the samples was assessed via HUMAnN pathway abundance analysis comparing M_abx and F_abx at 10 weeks of age. Further statistical validation of the pathways was performed using MaAsLin2. Upon analysis of the top 50 most abundant pathways in F_abx, several biosynthesis-related pathways were significantly enriched in abundance relative to M_abx (**Figure 3A**). These included amino acid biosynthesis (L-arginine, L-valine, L-ornithine, aromatic amino acid biosynthesis, tryptophan biosynthesis), nucleotide biosynthesis (5-aminoimidazole ribonucleotide biosynthesis), cell-structure biosynthesis pathways (peptidoglycan biosynthesis, UDP-N-acetylmuramoyl-pentapeptide biosynthesis), cofactor biosynthesis (folate transformations), carbohydrate biosynthesis (sucrose biosynthesis), carboxylate biosynthesis (chorismite biosynthesis), and lipid biosynthesis (fatty acid biosynthesis).

**Figure 3.**
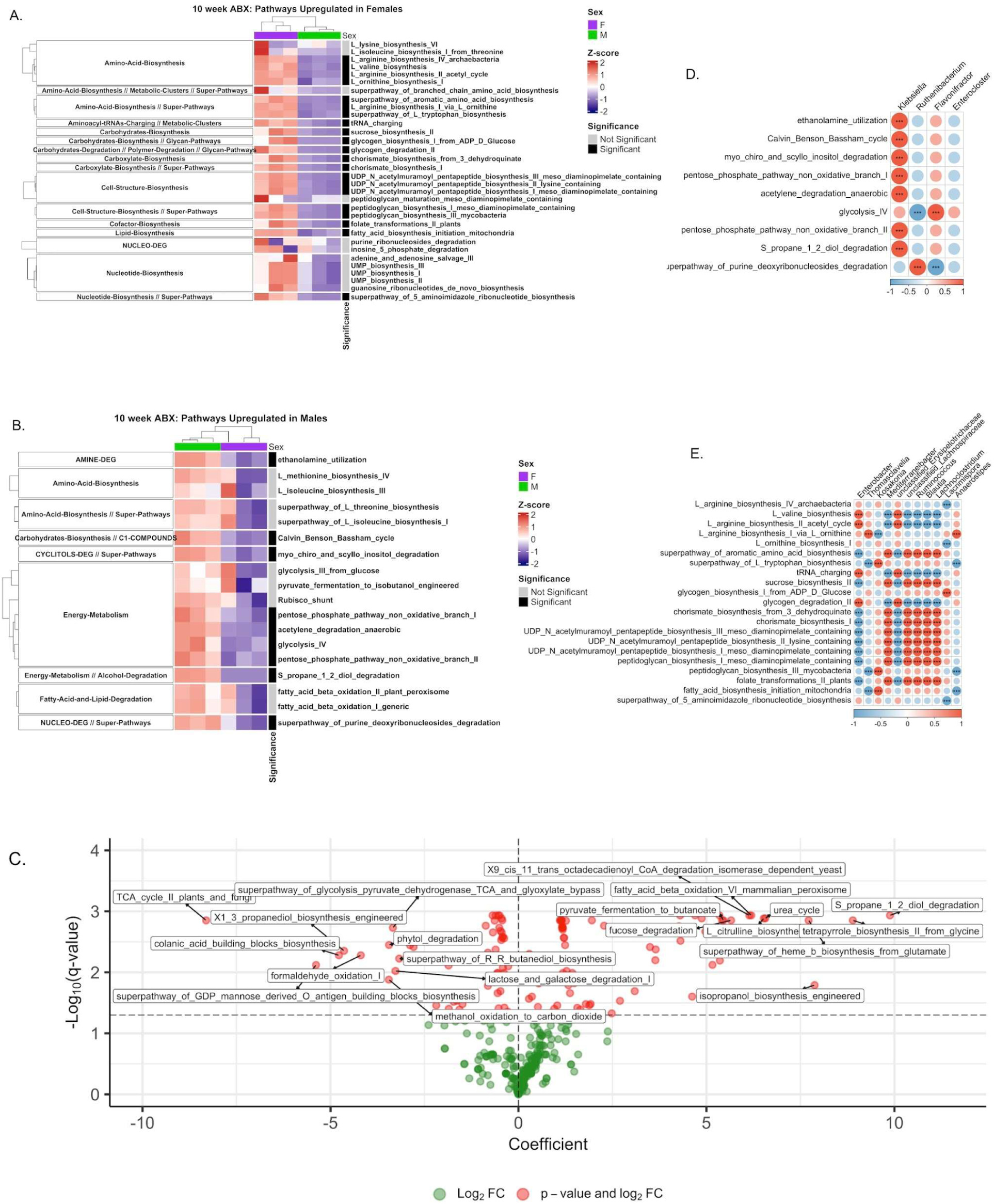
**(A)** Heatmap of functional pathways enriched in F_abx relative to M_abx at 10 weeks of age revealed F_abx microbiomes were enriched in biosynthesis-related pathways. **(B)** Heatmap of functional pathways enriched in M_abx relative to F_abx at 10 weeks of age revealed M_abx microbiomes were enriched in carbohydrate utilization, fermentation, energy metabolism, and substrate degradation pathways. **(C)** MaAsLin2 volcano plot of functional pathways between M_abx and F_abx at 10 weeks of age (enriched in M_abx top left, enriched in F_abx top right) confirmed enrichment of anabolic-driven processes in F_abx and enrichment of catabolic-driven processes in M_abx. **(D)** Spearman correlation analysis between differentially abundant taxa and pathways in M_abx at 10 weeks of age revealed many functional pathways were associated with *Klebsiella* (*** = BH-adjusted p-value < 0.001). **(E)** Spearman correlation analysis between differentially abundant taxa and pathways in F_abx at 10 weeks of age revealed functional pathways were associated with more diverse taxa than in M_abx (*** = BH-adjusted p-value < 0.001).

In comparison, M_abx samples at 10 weeks of age showed a significant enrichment of several pathways that were associated with energy metabolism-related activities (glycolysis, pentose phosphate pathways, anaerobic acetylene degradation, propane-1-2-diol degradation), nucleotide/nucleoside degradation (purine deoxyribonucleosides degradation), amine degradation (ethanolamine utilization), and cyclitol degradation (myo- chiro- scyllo- inositol degradation) (**Figure 3B**). An additional significant pathway included carbohydrate biosynthesis via the Calvin cycle, and though not statistically significant, fatty acid beta-oxidation pathways were also upregulated in M_abx. Overall, the pathway abundance analysis suggested that F_abx samples were enriched for anabolic and biomass-building functions, whereas M_abx samples were comparatively enriched for carbohydrate utilization, fermentation, energy metabolism, and substrate degradation pathways. MaAsLin2 analysis followed by visualization of the top 50 most abundant pathways was also performed to compare M_abx and F_abx at five weeks of age, and to evaluate the effect of antibiotics across different ages in M_abx and F_abx (**Supplemental Figures 17-19**).

The functional pathway divergence between M_abx and F_abx at 10 weeks of age was further visualized through a volcano plot of MaAsLin2 coefficients and q-values for all 371 pathway variables compared between M_abx and F_abx at 10 weeks of age. Among the most significant pathways enriched in females were the TCA cycle II (plants and fungi), superpathway of glycolysis-pyruvate dehydrogenase-TCA and glyoxylate bypass, colonic acid building block biosynthesis, and superpathway of GDP-mannose-derived O-antigen building block biosynthesis. Additional significant pathways in the volcano plot included 1,3-propanediol biosynthesis, lactose and galactose degradation I, formaldehyde oxidation I, methanol oxidation to carbon dioxide, and phytol degradation. In the M_abx mice, the urea cycle and S-propane-1,2-diol degradation, pyruvate fermentation to butanoate, L-citrulline biosynthesis, fucose degradation, tetrapyrrole biosynthesis II from glycine, and isopropanol biosynthesis were among the most significantly enriched pathways (**Figure 3C**). The overall distribution of the volcano plot demonstrated a broadly symmetric enrichment of significant pathways on both sides of the coefficient axis, indicating the functional metabolic divergence between M_abx and F_abx was not driven by enrichment in one sex alone but represented a bidirectional remodeling of the functional pathway landscape.

A Spearman correlation analysis was performed on selected abundant and significant pathways versus corresponding taxa in M_abx and F_abx. It was observed that *Klebsiella* was significantly correlated with several pathways in M_abx (**Figure 3D**), versus in F_abx the correlations were more distributed across multiple taxa and pathways, with commensals such as *Blautia*, *Ruminococcus*, *Lachnoclostridium*, *Mediterraneibacter*, and *unclassified Lachnospiraceae* showing coordinated correlations across several biosynthesis pathways (**Figure 3E**).

To further study the effects of antibiotic perturbation on microbial metabolism, untargeted metabolomics data from the cecal contents of the same mice was analyzed. Principal Component Analysis (PCA) on all metabolomics samples demonstrated a clear treatment-based separation along the first component followed by an age specific separation along the second component (**Supplemental Figure 20**). To further investigate the samples at 10 weeks of age, an UpSet plot was used to visualize the overlap of detected metabolite features across all groups at 10 weeks of age (**Figure 4A**). The largest intersection consisted of 1803 features, which were metabolites shared across all groups, demonstrating that the core metabolome detected was relatively common across all experimental conditions. 215 features were shared between M_abx and F_abx and 106 features were shared between M_control and F_control demonstrating treatment specific differences. 53 metabolites were present in F_abx only and nine metabolites were present in M_abx only, highlighting sex-specific differences in metabolite presence within the treatment groups.

**Figure 4.**
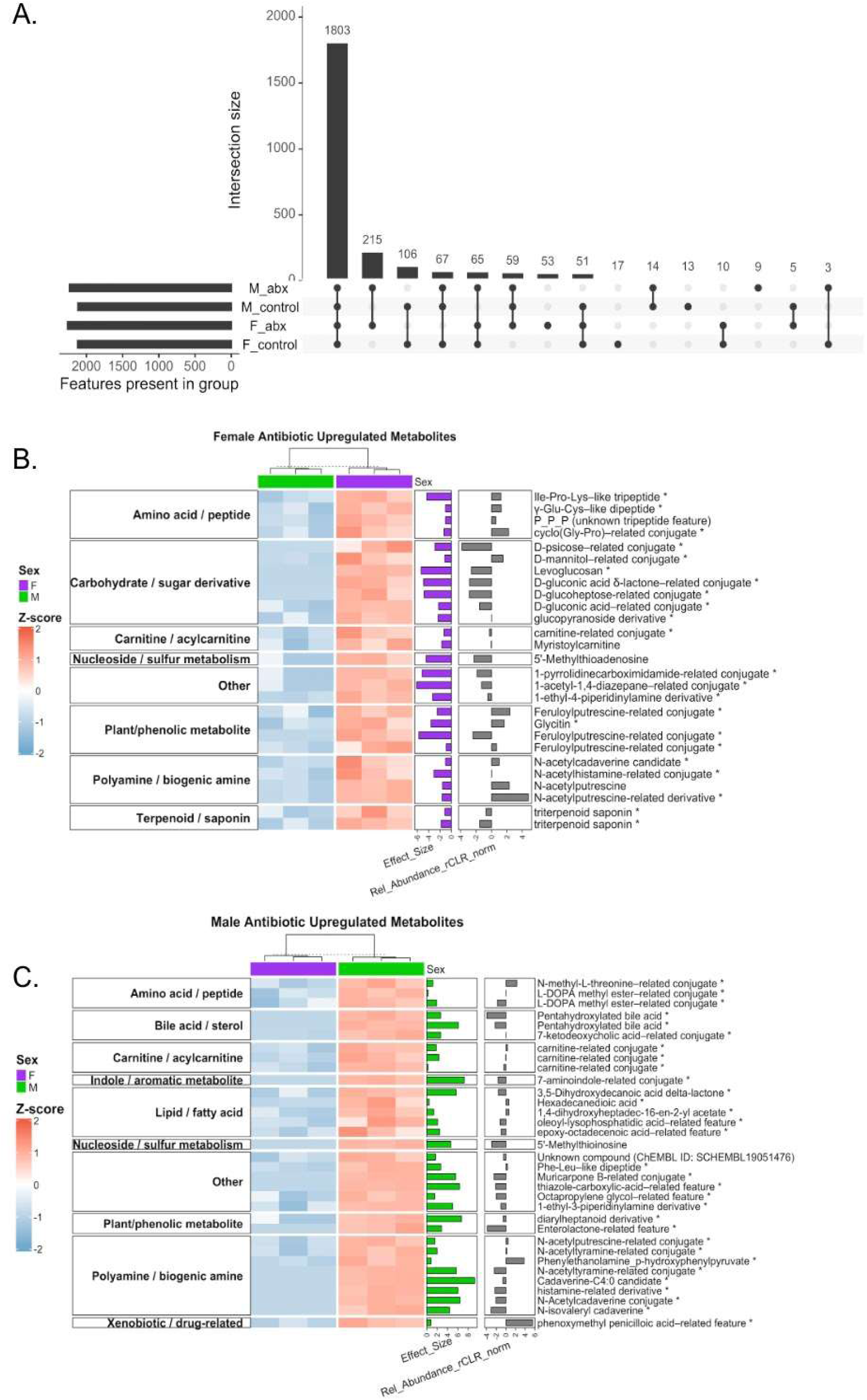
**(A)** UpSet plot summarizing metabolite presence across groups at 10 weeks of age revealed subsets of metabolites potentially associated with antibiotic-induced and sex-specific metabolic responses. **(B)** Heatmap of cecal metabolites significantly enriched in F_abx relative to M_abx at 10 weeks of age. **(C)** Heatmap of cecal metabolites significantly enriched in M_abx relative to F_abx at 10 weeks of age. For both heatmaps (B-C) only metabolites identified as significant by linear regression analysis (BH-adjusted p < 0.05) that possessed compound name annotations were included in the visualizations.

To further characterize sex-differences in the antibiotic group at 10 weeks of age, a partial least squares discriminant analysis (PLS-DA) was performed and features with a VIP > 1 and a |Cliff’s delta| > 0.5 were subject to linear regression using the lm() feature in R. To validate the results of the PLS-DA analysis, permutation testing was performed. The observed balanced error rate (PER = 0.167) was significantly lower than expected under random class assignment (permutation p = 0.004), indicating that the model’s classification performance was unlikely to have occurred by chance and reflects meaningful separation between the groups (**Supplemental Figure 21**). From the linear regression analysis, 60 differentially abundant annotated metabolites between M_abx and F_abx mice were present with an adjusted p-value < 0.05. Among these 60 significant annotated metabolites, 27 were upregulated in females and 33 were upregulated in males, revealing sex-specific differences in metabolite profiles.

Notably, F_abx was enriched in several carbohydrate and sugar derivatives, including D-psicose-related-conjugate, D-mannitol-related-conjugate, levoglucosan, D-gluconic acid-related features, D-gluconic acid δ-lactone-related conjugates, D-glucoheptose-related conjugates, and glycopyranoside derivatives (**Figure 4B**). F_abx also showed enrichment of multiple peptide and amino acid-associated metabolites, including γ-Glu-Cys-like dipeptides, cyclo(Gly-Pro)-related conjugates, Ile-Pro-Lys-like tripeptides, and plant/phenolic-associated metabolites such as glycitin, feruloylputrescine-related conjugates. Lastly, a significant enrichment and abundance of biogenic amines such as several N-acetylputrescine derivatives, along with an N-acetylcadaverine and N-acetyl histamine derivative were observed in F_abx.

In contrast, M_abx samples predominantly demonstrated enrichment of polyamines and biogenic amines, specifically N-acetylcadaverine, cadaverine-associated features, N-acetyltyramine-related conjugates, histamine-related derivatives, N-acetylhistamine-related conjugates, as well as an N-acetylputrescine derivate (**Figure 4C**). Males also exhibited enrichment of bile acid-associated metabolites, including a pentahydroxylated bile acid feature and a 7-ketodeoxycholic acid-related conjugate and lipid-associated metabolites such as hexadecanedioic acid, oleoyl-lysophosphatidic acid-related features, epoxy-octadecenoic acid-related features, and 3,5-dihydroxydecanoic acid delta-lactone. Additional male-enriched metabolites included an indole/aromatic-associate 7-amino-indole conjugate, xenobiotic/drug-related metabolite phenoxymethyl penicilloic acid, and several compounds classified as unknown. Collectively, these results demonstrated a sex-based separation in metabolite composition following antibiotic treatment. Spearman correlation analyses were performed for corresponding M_abx and F_abx metabolites against their corresponding microbial pathways and taxa. However, the results did not reveal clearly biologically interpretable correlation patterns (**Supplemental Figures 22-25**).

### Multi-omics integration revealed antibiotic-induced sex differences were detectable in splenic transcriptomics

To study the structure of variation in splenic transcriptomics across all groups at 10 weeks of age, principal component analysis (PCA) was performed on variance-stabilizing transformed (VST) count data from all samples (**Figure 5A**). The first principal component (PC1) explained 39% of the variance, whereas the second principal component (PC2) explained 25% of the variance. Notably, the samples were separated by sex and when looking at the female samples (F_abx and F_control) there was a greater separation along PC1 and PC2 relative to the separation observed in male samples (M_abx and M_control). This observation suggested that antibiotic treatment was likely associated with a larger transcriptomic shift in the spleens of female mice relative to male mice.

**Figure 5.**
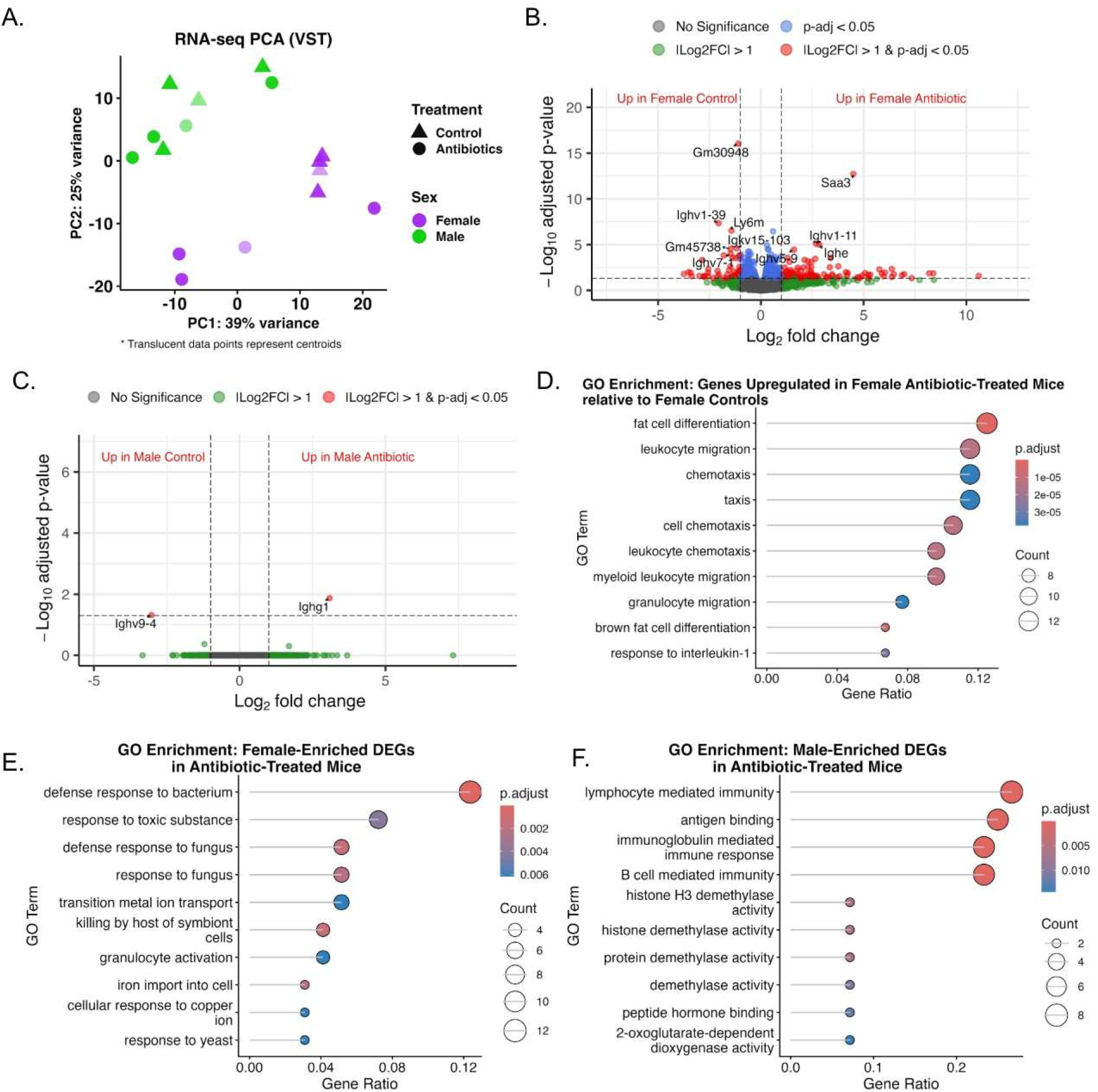
**(A)** PCA of variance-stabilized transformed (VST) RNAseq expression data at 10 weeks of age revealed a significant effect of sex (R^2^ = 0.177, p = 0.022), but not treatment (R^2^ = 0.106, p = 0.211). Within-sex analyses showed no significant treatment effect in females (R² = 0.273, p = 0.20) or males (R² = 0.144, p = 0.80) **(B)** Volcano plot of differential gene expression between F_abx and F_control at 10 weeks of age revealed a robust transcriptional response to antibiotic exposure. **(C)** Volcano plot of differential gene expression between M_abx and M_control at 10 weeks of age revealed an attenuated transcriptional response to antibiotic exposure relative to females. **(D)** Gene ontology (GO) enrichment analysis of genes significantly upregulated in F_abx relative to F_control at 10 weeks of age revealed a directional trafficking of innate immune cells. **(E)** GO enrichment analysis of genes significantly upregulated in F_abx relative to M_abx at 10 weeks of age revealed a transcriptional response associated with innate immune defense and microbial recognition. **(F)** GO enrichment analysis of genes significantly upregulated in M_abx relative to F_abx at 10 weeks of age revealed a transcriptional response associated with adaptive immunity and antibody mediated processes.

To characterize the effect of antibiotic treatment on splenic gene expression in each sex, DESeq2 differential analysis was performed using a ∼Sex x Treatment interaction design. In female mice, the antibiotic treatment was associated with a substantially greater number of differentially expressed genes (DEGs) than in males. The female volcano plots revealed a large number of genes meeting the significance threshold (**Figure 5B**). Specifically, F_abx had significantly upregulated genes such as *Saa3* (serum amyloid A3), *Ighv1-11, Ighv5-9*, and *Ighe*, among many others, several of which were associated with immunoglobulin variable or constant region segments (*Igh*), which suggested antibiotic associated changes in B-cell or antibody-related gene expression. Moreover, *Saa3* is an acute phase reactant, whole upregulation is associated with inflammatory conditions and innate immune activation ^40^.

In contrast, the male volcano plot revealed a drastically attenuated transcriptomic response to antibiotic treatment (**Figure 5C**). Specifically, only two genes reached significance in M_abx vs. M_control analysis. *Ighg1*, encoding the *IgG1* heavy chain constant region was significantly upregulated while *Ighv9-4* was significantly downregulated. The result that only two DEGs reached significance, provided further evidence that antibiotic treatment had a minimal detectable effect on global splenic gene expression in male mice, as compared to a more extensive difference observed in females.

To characterize the biological processes associated with the genes upregulated in F_abx relative to F_control, Gene Ontology (GO) enrichment analysis was performed using the clusterProfiler package (**Figure 5D**). Among the top enriched processes in F_abx, there was fat cell differentiation (with the highest gene ratio of 0.13), migration (leukocyte, myeloid leukocyte, granulocyte), chemotaxis (cell, leukocyte), brown fat cell differentiation, and response to interleukin-1. The predominance of leukocyte migration, chemotaxis, and granulocyte-related terms among the top enriched processes suggests that the upregulated genes were potentially involved in the directional trafficking of innate immune cells such as granulocytes and myeloid leukocytes.

Given that only *Ighg1* was significantly upregulated in M_abx relative to M_control, a GO enrichment analysis did not yield meaningful interpretable results (**Supplemental Figure 26**). However, *Ighg1* codes for the protein that enables antigen binding activity and is involved in B cell differentiation, suggesting potential regulation of an adaptive immune response ^41^.

To directly compare splenic gene expression between M_abx and F_abx, DESeq2 was performed on the antibiotic group alone using ∼Sex as the design (**Supplemental Figure 27**). The sex-specific DEG differences were heavily associated with X-linked and Y-linked genes such as *Xist* in females and *Kdm5d, Uty*, *Ddx3y*, and *Eif2s3y* in males respectively. These genes were also differentially expressed in a comparison of M_control vs. F_control (**Supplemental Figure 28**). This is in line with biological sex being an inherently dominant source of global transcriptomics variance in male and female spleens and may also explain why a dominant sex-based separation was seen in the PCA plot ^42^. Although the presence of DEGs in F_abx such as *Saa3*, *Ngp*, and *Ltf* and *Ly6m* and several *Igh-*related genes in M_abx suggested that the transcriptomic state of antibiotic-treated male and female spleens differed not only in the magnitude of the treatment response relative to their controls but also different from each other, with females showing elevated acute-phase and innate immune-associated transcripts ^40^.

To characterize the biological processes associated with the sex-specific DEGs identified in the antibiotic group, GO enrichment analysis was performed separately on F_abx and M_abx enriched DEGs from the ∼Sex model. From the genes significantly upregulated in F_abx, the GO terms were centered on innate immune defense and microbial recognition with top terms being related to responses to bacterium, fungus, toxic substance, and yeast as well as transition metal ion transport, iron import into cell, cellular response to copper iron, granulocyte activation, and killing by host of symbiont cells (**Figure 5E**).

M_abx had upregulated DEGs that linked to GO terms associated with adaptive immunity and antibody mediated processes (**Figure 5F**). This included lymphocyte mediated immunity, antigen binding, immunoglobulin mediated immune response, and B-cell mediated immune response, which were the four terms with the highest gene ratio. Additional terms included H3 demethylase activity, histone demethylase activity, protein demethylase activity, demethylase activity, peptide hormone binding, and 2-oxoglutarate-dependent dioxygenase activity. These pathways may represent epigenetic regulatory differences in males relative to females, with some pathways likely being associated with the upregulation of Y-linked genes such as *Kdm5d* that is linked to histone demethylase activities ^43^. Overall, the breadth of the terms in F_abx suggested a broad innate immune defense gene enrichment relative to M_abx.

To evaluate whether the sex-specific antibiotic associated differences that we identified across individual omics layers converged into coherent multi-omics signatures, an integrative analysis was performed using DIABLO implemented in the mixOmics package in R. The model integrated five matched omics blocks (taxa metagenomics, ARG metagenomics, metagenomic functional pathways, untargeted metabolomics, and bulk RNA transcriptomics). The model was run with a design matrix value of 0.5, two latent components (Comp1 and Comp2), and a sparse feature selection retaining ten features per block per component. The final model was validated using permutation testing and it demonstrated that the addition of a second latent component decreased the error rate in the model (**Supplemental Figure 29**).

Individual block sPLS-DA scatter plots demonstrated the degree to which each omics block discriminated against the four experimental groups on the two components (**Figures 6A, Supplemental Figures 30-33A**). Consistently in all five blocks F_abx and M_abx samples separated from their respective controls. Control samples formed intermediate clusters that were not strongly separated from each other in any block, suggesting that the dominant discriminative signal in the integrated model was antibiotic treatment. Across all blocks, F_abx and M_abx separated across both Comp1 and Comp2. F_abx separated from the controls along Comp1, whereas M_abx separated from the controls along Comp2. The taxa block provided the strongest discrimination on Comp1 with 55% explained variance and 24% on Comp2. The RNAseq block demonstrated the weakest overall separation of the five blocks on both components.

**Figure 6.**
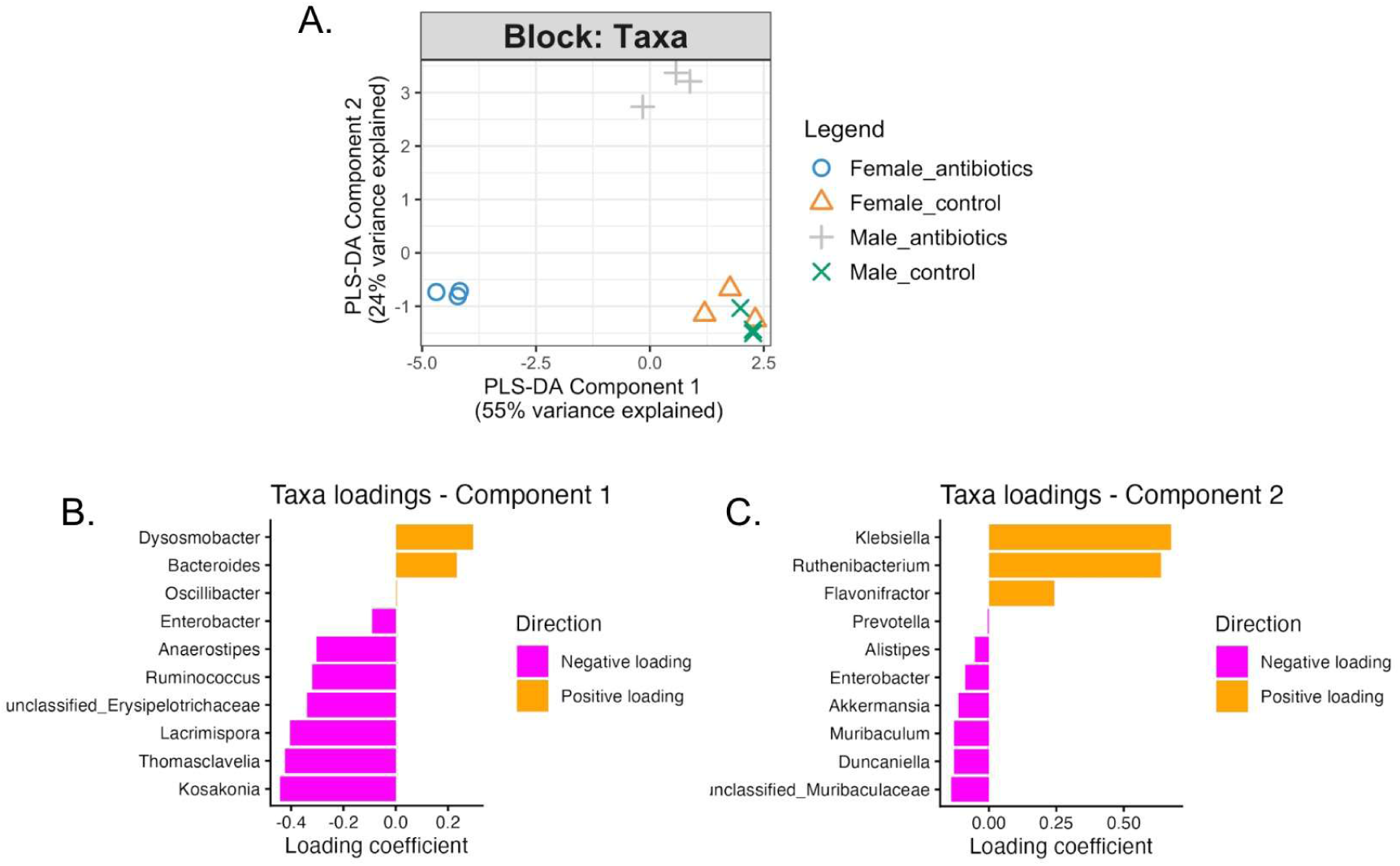
**(A)** PLS-DA individual block plot from DIABLO showed sample separation based on taxonomic composition at 10 weeks of age. **(B-C)** Loading plots showed the 10 taxa with the largest absolute loading coefficients contributing to Components 1 and 2, respectively.

In the taxa block, features that contributed to Comp1 were *Dysosmobacter*, *Bacteroides*, *Oscillobacter* with a positive coefficient and *Enterobacter*, *Anaerostipes*, *Ruminococcus*, *Lacrimispora*, *Thomasclavelia*, *Kosakonia*, and *Erysipelotrichaceae* with a negative coefficient. These negative taxa loadings were in agreement with previously observed taxa that were differentially abundant in F_abx (**Figure 6B**). Along Comp2, *Klebsiella*, *Ruthenibacterium*, and *Flavonifractor* had positive loadings in agreement with taxa that were differentially abundant in M_abx (which separated from F_abx and controls along Comp2) (**Figure 6C**).

In the ARG block, along Comp1 most selected features loaded negatively, including several genes associated with multidrug efflux pumps that were differentially abundant in F_abx (**Supplemental Figure 30B**). Along Comp2, all ten selected ARGs loaded positively suggesting upregulation in M_abx (**Supplemental Figure 30C**). In the metagenomics functional pathway block Comp1, the positive loadings were largely comprised of fermentation pathways (pyruvate to propanoate I, acetyl-CoA to butanoate II), whereas the negative loadings were largely comprised of biosynthesis (molybdopterin, 1,3-propanediol biosynthesis, superpathway of 2,3-butanediol) pathways (**Supplemental Figure 31B**). On Comp2, positive loadings were associated with redox metabolism and negative loadings included several biosynthesis (peptidoglycan III, chorismate I, dTDP-β-L-rhamnose) and degradation (guanosine nucleotides I, D-arabinose I) pathways (**Supplemental Figure 31C**). In the metabolomics block, Comp1 had positively loaded metabolite features that suggested increased host processing of microbial products. Negatively loaded metabolite features included those that were associated with polyamine and nitrogen metabolism (**Supplemental Figure 32B**). Along Comp2, positive metabolite features included those that were associated with lipid remodeling, oxidative metabolism, and protein and amino acid turnover. Negative loadings included metabolites associated with metabolic stress (**Supplemental Figure 32C**). In the bulk RNA transcriptomics block Comp1, positive genes included those associated with lymphocyte and macrophage function (**Supplemental Figure 33B**). On Comp2, positive features consisted mostly of uncharacterized transcripts and negative features included those associated with cell-surface signaling proteins in immune cell communication (**Supplemental Figure 33C**).

A Clustered Image Map (CIM) was generated and within the M_abx cluster, several correlated features across all five omics datasets were hierarchically clustered together (**Figure 7**). A distinct cluster of upregulated features were observed in M_abx and F_abx, which suggested the existence of a correlation between the five omics datasets in a sex-specific manner following antibiotic exposure. Interestingly, two more clusters were observed in the CIM which demonstrated a selective downregulation in F_abx and M_abx and moderate to high expression in other groups. The formation of four distinct clusters was further validated by the correlation circle plot (**Supplemental Figure 34**).

**Figure 7.**
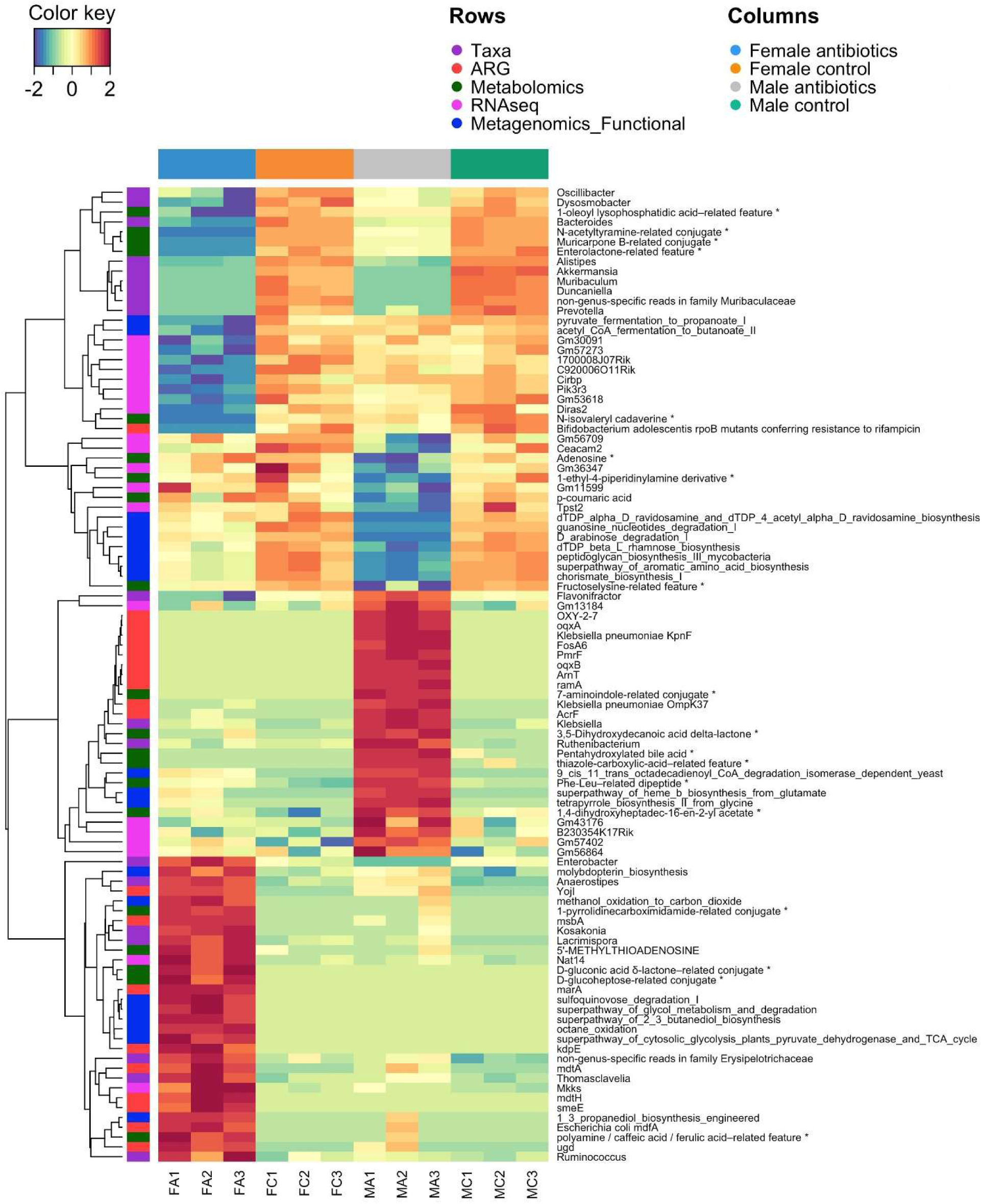
Clustered Image Map (CIM) of features selected by the DIABLO model at 10 weeks of age.

To improve cluster visualization, Ward’s minimum variance method was applied to the DIABLO CIM with a cluster value of four. Among the four clusters that were generated (**Supplemental Figure 35**), one Ward cluster of features was upregulated in F_abx (Cluster 1), one upregulated in M_abx (Cluster 2) and two were downregulated in the Antibiotic groups and upregulated in the Control groups (Cluster 3 and 4). Cluster 3 was more downregulated in the M_abx group relative to F_abx in which mean expression levels were marginally over 0. Cluster 4 was more downregulated in F_abx relative to M_abx in which mean expression levels were marginally below 0. Dendrogram plots were generated for each cluster and within Cluster 1, correlated features across all five omics datasets hierarchically clustered together (**Figure 8A**). Specifically, this included seven taxa (*Enterobacter*, *Anaerostipes*, *Kosakonia*, *Lacrimispora*, *Thomasclavelia*, *Ruminococcus*, *Erysipelotrichaceae*), nine ARGs, eight functional pathways, five metabolites, and two RNAseq genes. The results from the Cluster 2 dendrogram specifically showed the three M_abx enriched taxa *Flavonifractor*, *Klebsiella*, and *Ruthenibacterium* along with 10 ARGs, three metagenomic functional pathways, six metabolites, and four RNAseq genes (**Figure 8B**). The third cluster was characterized by increased suppression in M_abx and was comprised of seven functional pathway features, five RNAseq genes, and four metabolite features (**Figure 8C**). Finally, the dendrogram of the fourth cluster included features that had increased suppression in F_abx samples relative to M_abx, while control groups maintained elevated abundance (**Figure 8D**). The suppression specific clusters across the five omics layers in M_abx and F_abx represented an additional dimension of the sex-dependent antibiotic response captured by the DIABLO integration.

**Figure 8.**
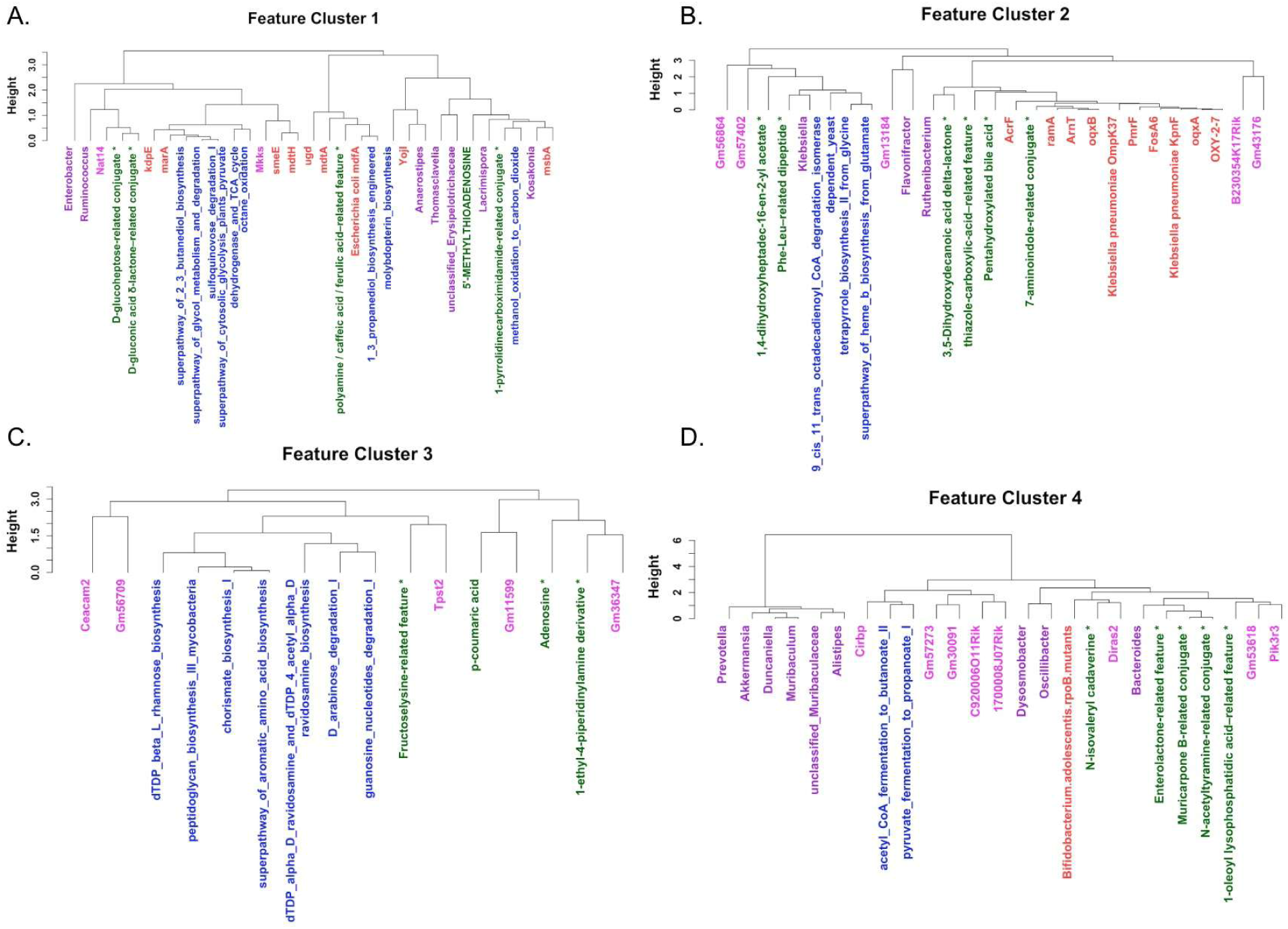
Dendrogram of features assigned to Cluster 1 following Ward hierarchical clustering of DIABLO-selected features **(A)** enriched in F_abx, **(B)** enriched in M_abx, **(C)** suppressed in M_abx, and **(D)** suppressed in F_abx.

## Discussion

The objective of this study was to decipher the effect of antibiotics on the gut microbiome across sexual maturity in a sex-stratified manner. The results of this study revealed that chronic antibiotic exposure across sexual maturation reshaped the gut microbiome and splenic gene expression in a fundamentally sex-specific manner.

A prominent indication of a sex-specific divergence was in the alpha diversity at 10 weeks of age in which F_abx mice exhibited significantly higher Shannon and Simpson diversity indices compared to M_abx mice. This finding was consistent with a growing body of evidence demonstrating that biological sex shapes the composition, diversity, and resilience of the gut microbiome ^15^. Moreover, after one week of antibiotic exposure, there was a significant drop in alpha diversity in both M_abx and F_abx at five weeks of age with both sexes unanimously dominated by *Kosakonia*. The near-uniform dominance by *Kosakonia* likely reflected the collapse of colonization resistance rather than a host-sex interaction. The opportunistic bloom of *Kosakonia* was consistent with its capacity as a facultative anaerobe to exploit the oxidative and nutrient-rich niche depletion of the obligate anaerobic community ^44^. The absence of a sex-difference prior to sexual maturity was consistent with previous observations that sex differences in the microbiome generally emerge after puberty ^15,22,23^. F_abx samples showed higher diversity at 10 weeks of age and the distribution of microbes spanned both opportunistic *Enterobacteriaceae* (*Enterobacter*, *Kosakonia*) and commensal-associated taxa (*Blautia*, *Ruminococcus*, *Lachnoclostridium*, *Lacrimispora*, and *Mediterraneibacter*) that were also present in the control samples ^45^. The partial re-emergence of these microbes in females under sustained antibiotic pressure may be indicative of sex-specific differences in susceptibility to prolonged antibiotic-associated microbial depletion. A plausible explanation for the diversification observed in females may be related to the influence of sex hormones on the microbiome. At 10 weeks of age, C57Bl/6 mice are expected to have reached sexual maturity, with females exhibiting elevated estrogen levels relative to pre-pubertal stages ^46^. Estrogen has been proposed to exert a positive regulatory influence on gut microbial diversity, with low estrogen states associated with decreased diversity and dysbiosis ^47,48^.

M_abx were dominated by low microbial diversity at 10 weeks of age, primarily composed of *Klebsiella*, *Flavonifractor*, and *Ruthenibacterium*. The near-complete elimination of *Kosakonia* (dominant genera at five weeks of age) and the failure to diversify towards commensal-associated taxa suggested that the M_abx gut remained in a persistently dysbiotic state. This was consistent with previous observations that the response of the rodent gut microbiome to antibiotics differs between males and females ^6,49^. The enrichment of *Klebsiella oxytoca* in M_abx mice at 10 weeks of age was a notable finding of this study as *K. oxytoca* has been documented as a causative organism of antibiotic associated hemorrhagic colitis ^50^. Though not formally quantified in this study, a notably enlarged ceca in M_abx were observed relative to F_abx mice showing consistency with *Klebsiella-associated* intestinal pathophysiology, as cecal enlargement can reflect antibiotic-induced metabolic waste accumulation and a shift toward a pathogenic microbial environment ^50^.

An intriguing question arises as to why *Klebsiella* expanded in males and not in females. While future studies are needed for a definitive answer, this study offers plausible explanations upon integration of resistome, microbial functional pathways and metabolomics data. The co-enrichment of the *OXY-2-7* ARG with *Klebsiella* in M_abx was mechanistically coherent because *K. oxytoca* possesses intrinsic OXY-type beta-lactamases as part of its core genome, meaning that ampicillin exposure could selectively favor the survival and proliferation of *Klebsiella* at the expense of susceptible commensal organisms ^51^. The observation that a putative (phenoxymethyl)penicilloic acid metabolite showed significantly higher abundance in M_abx relative to F_abx at 10 weeks of age was consistent with higher beta-lactamase activity in males (present with elevated *OXY-2-7*) because penicilloic acids are formed through enzymatic hydrolysis of the beta-lactam ring by beta-lactamases. The combination of *Klebsiella* expansion, *OXY-2-7* enrichment, and enrichment of putative penicilloic acid provide a multi-layered picture of antibiotic driven selective pressure in M_abx mice that was not observed in F_abx.

In F_abx mice, the resistome at 10 weeks was characterized by the re-emergence of tetracycline resistance genes (*tet(40)*, *tet(O)*). These ARGs were significantly correlated with several commensal-like microbes in F_abx. Tetracycline resistance genes are among the most widely present ARGs in gut microbiomes ^52^. The re-emergence of commensal-associated taxa in F_abx suggested that the female resistome at 10 weeks of age exhibited a more distributed ARG architecture, rather than the opportunistic-driven resistance profile observed in males.

This sex difference was further supported by a functional pathway divergence of the metabolic landscape of M_abx and F_abx communities at 10 weeks of age. F_abx samples had a significant enrichment in several biosynthesis-oriented pathways. This anabolic pathway profile was consistent with communities that are engaged in active biomass production and biosynthetic investment, potentially reflective of a metabolically diverse and expanding microbial ecosystem. In contrast, M_abx samples showed a significant enrichment of pathways dominated by energy dissipation, substrate catabolism, and fermentation. This catabolic, fermentation-heavy profile suggested a community that was prioritizing rapid energy acquisition rather than biosynthetic investment. We hypothesized that this was reflective of the strategy of fast-growing opportunistic organisms that were exploiting a low-competition, substrate-rich environment after the collapse of the resident microbiota. Notably, the ethanolamine utilization pathway was significantly enriched in M_abx relative to F_abx. Ethanolamine is abundant under dysbiotic conditions and can be metabolized as a carbon and nitrogen source providing a colonization advantage to opportunistic pathogens, such as *Klebsiella* ^53,54^. The enrichment of this pathway in M_abx may therefore represent *Klebsiella* exploiting host-derived substrates that became available in the M_abx environment, potentially contributing to its success in colonization.

The untargeted cecal metabolomics data provided complementary evidence that M_abx and F_abx communities functionally diverged as reflected in their distinct profiles. Both M_abx and F_abx samples were significantly enriched for putative polyamine and biogenic amine-associated metabolites (cadaverine, putrescine, histamine) relative to controls. Notably, *K. oxytoca* has been shown to produce cadaverine via decarboxylation of lysine ^55,56^. The enrichment of biogenic amines in M_abx, alongside the taxonomic dominance of *Klebsiella* and the metabolic pathway signatures of catabolic fermentation provide a systems-level insight into mechanisms associated with dysbiotic gut communities. F_abx on the other hand showed relatively higher levels of putrescine-related amines. *Enterobacter* has been shown to use putrescine for acid resistance, biofilm formation, and modulation of environmental stress ^57,58^. Putrescine supplementation has also been shown to increase alpha diversity of the gut microbiome and reduce inflammation, providing some mechanistic insight into the more resilient dysbiotic state in F_abx ^58^. Further, in F_abx at 10 weeks of age, glycitin was significantly upregulated. Glycitin, a phytoestrogen that can interact with estrogen receptors, is subject to microbial biotransformation by isoflavone-metabolizing *Ruminococcus*, a genera that emerged in F_abx at 10 weeks of age ^59^. The differential enrichment of glycitin in F_abx raises the possibility that microbiome-mediated phytoestrogen metabolism may influence the relationship between estrogen and the microbiome^60^.

The splenic bulk RNAseq data revealed a striking asymmetry in the gene expression between M_abx and F_abx. There was a clear sex-based separation in the PCA plot, likely related to the strong expression of X-linked *Xist* in females and Y-linked genes *Kdm5d*, *Uty*, *Ddx3y*, and *Eif2s3y* in males. F_abx showed a larger separation in the PCA plot and higher number of DEGs relative to its F_control, which suggested that beyond an expected sex-chromosome driven difference in transcription, the antibiotic treatment itself produced a larger transcriptional perturbation in female spleens compared to male. Only two DEGs were significant in M_abx relative to M_control, which suggested that either transcriptional response was less perturbed than in females or perturbed but failed to reach significance. From the present data it is not possible to resolve whether the larger transcriptomic shift in females reflected a more reactive immune system or greater sensitivity to gut-derived signals under antibiotic pressure. F_abx was functionally enriched in innate immune-associated pathways relative to F_control, which suggested a heightened innate inflammatory signaling. This was also supported by the upregulation of the *Saa3* gene, an acute phase reactant induced downstream of innate immune pathways ^61^. Collectively, these findings suggested that F_abx mice experienced a broader immune activation composed of both innate and adaptive components. In contrast, the only significantly upregulated gene in M_abx relative to M_control was *Ighg1*, encoding the IgG1 heavy chain constant region associated with B-cell activation and antibody production ^41^. This suggested that antibiotic exposure did induce a measurable humoral immune response in males, although the response appeared considerably narrower than the immune transcriptional remodeling observed in females.

This pattern of broader innate and adaptive immune-associated transcriptional responses in F_abx combined with a comparatively humoral-skewed response in M_abx, was further supported by the sex-stratified GO enrichment analysis in the antibiotic groups. Male-enriched DEGs were primarily associated with adaptive immune processes (lymphocyte-mediated immunity, antigen binding, immunoglobulin-mediated immune response, B-cell mediated immunity), consistent with expression of immunoglobulin-associated genes *Ighg1* and *Ighv1-31*. In contrast, female-enriched DEGs were associated with broader host defense and innate immune-associated functions (granulocyte activation, defense responses to bacteria and fungi), supported by upregulated innate defense genes *Saa3*, *Prtn3*, *Ngp*, *Chil3*, and *Ltf* ^40,62–64^. The enrichment of histone demethylase-related terms in males was likely driven by Y-linked genes (*Kdm5d* and *Uty*) which may reflect constitutive chromosomal sex differences rather than antibiotic-responsive immune remodeling^65,66^.

It is worth considering whether the post-pubertal hormonal environment may be associated with the sexually dimorphic splenic transcriptomics response and the differential immune landscape. While circulating hormone levels were not measured in this study, a substantial body of evidence supports the concept of estrogen being a plausible modulator of the immune phenotypes and raises the possibility to generate hypotheses that may help to further elucidate the observed sex differences^67–69^. F_abx exhibited enrichment of DEGs associated with granulocyte activation and leukocyte chemotaxis and estrogen receptors have been shown to be expressed on a range of innate immune cells (macrophages, neutrophils, etc.) and modulate leukocyte recruitment^70^. Beyond its effects on innate immunity, estrogen is a well-established regulator of adaptive immune responses, particularly B-cell activation and antibody production ^70,71^. This is consistent with the observation of F_abx demonstrating simultaneous upregulation of multiple immunoglobulin variable region genes along with enrichment of adaptive immune pathways. In contrast, M_abx did not exhibit the same broad innate transcriptional activation, but enrichment of more focused adaptive immunoglobulin-mediated processes. The male hormonal environment shapes immune responses differently and androgens such as testosterone are generally associated with altered immune polarization and differences in innate immune activation ^71^. It is important to note that the relationship between sex hormones and the gut microbiome is generally considered bidirectional: host hormones influence microbial composition and immune status, while the microbiome contributes to hormone metabolism and signaling ^72,73^. As such, the sex-specific immune responses observed here likely reflect dynamic interactions between a post-pubertal endocrine state, antibiotic-driven microbial disruption, and host immune programming rather than a hormonal effect alone ^71–73^.

The multi-omics integration produced meaningful exploratory results and many of the features selected and correlated through DIABLO recapitulated the sex-specific patterns identified in the independent analyses, particularly within taxa, ARGs, metagenomic functional pathways, and metabolomics. In contrast, RNAseq contributed comparatively less to the group discrimination across the latent components. This was reflected in the total explained variance from the PLS-DA DIABLO model, in which taxa accounted for the highest explained variance across the first two components (79%), followed by metagenomic functional pathways (75%), ARGs (73%), metabolomics (53%), and RNAseq explaining comparatively less (28%). Collectively, these results suggested that RNAseq represented the weakest contributor to group separation across the integrated datasets. Therefore, the RNAseq genes identified in the correlated multi-omics clusters (many of which remain poorly characterized), should be interpreted cautiously.

From the CIM and Ward’s cluster analysis, M_abx were predominantly enriched in *Klebsiella*, *Ruthenibacterium*, and *Flavonifractor* along with multiple ARGs. The metabolites and metagenomic functional pathways in this cluster were also previously identified as significantly upregulated in M_abx and were related to tetrapyrrole biosynthesis and fatty acid/lipid degradation. Metabolites identified in this cluster at a high-level were classified as bile acids, indole, lipid/fatty acids, and amino acid/peptides, which suggested an altered bile acid and aromatic metabolite metabolism with antibiotic perturbation. The cluster enriched in F_abx was characterized by co-enrichment of a more taxonomically diverse microbial community along with ARGs that were associated with the sex-specific resistome patterns that were previously identified in the study. Functionally, this cluster was associated with pathways that are involved in energy generation, alcohol and carbohydrate degradation, and cofactor and vitamin biosynthesis. While some of these pathways were not individually significant in the earlier MaAsLin2 analyses, their appearance in the integrated cluster was notable. Specifically, while the most abundant and significant microbial functional pathways from the MaAsLin2 analysis were largely biosynthesis-oriented, this cluster incorporated both biosynthetic and degradative pathways, potentially reflecting a metabolically adaptable microbial community. This interpretation was further supported by the metabolite composition of the cluster that was enriched in carbohydrate-derived metabolites and sulfur metabolism-associated compounds, suggesting active carbon use and metabolic turnover. Interestingly, metagenomic functional pathways contributed comparatively less explained variance along component 1 (the primary axis separating F_abx samples) relative to the stronger contribution observed along component 2 in males. This may partly explain why several pathways that emerged in the F_abx cluster differed from those highlighted in the independent differential analyses.

Among the two clusters that showed suppressed activity in the antibiotic treated groups, the cluster with higher suppression in F_abx was characterized by reduced abundance of microbial fermentation-associated pathways along with depletion of secondary metabolites. In contrast, the cluster that showed increased suppression in M_abx consisted of microbial biosynthetic pathways in aromatic amino acid metabolism, nucleotide turnover, and structural carbohydrate synthesis, potentially reflective of reduced biosynthesis activity under male-associated dysbiosis. However, the biological significance of these suppression patterns remains exploratory and warrants further investigation.

Several limitations need to be considered for the present study. First, the exploratory nature of the work and the small sample size (n=3 per group) limited statistical power, and therefore, the findings from this study should be interpreted as hypothesis-generating to identify candidate microbiome changes associated with sex rather than conclusive. Due to the exploratory nature of this work, mice were housed according to sex and treatment groups with one cage per group. This partially confounded cage- and treatment-effects on the microbiome due to the coprophagic nature of mice. Follow-up replication studies using multiple cages per treatment group will help to disentangle this as a confounding factor. Nevertheless, the longitudinal microbiome data collected in this study, particularly in the Control groups, demonstrated the consistency in the microbiome across cages and the drastic and dynamic effect of the antibiotic perturbation on microbial community composition. The untargeted metabolomics data also suffered from incomplete annotation, meaning that a significant proportion of differentially abundant features remained uncharacterized and their biological relevance could not be assessed. Further, the compositional microbiome data that was measured as relative abundance only reflected proportional shifts in a closed ecosystem and did not capture absolute microbial abundances, which further limited mechanistic interpretation. Microbial pathways that were interpreted from shotgun metagenomics data could only infer the metabolic potential of the community and could not capture actively transcribing pathways. In terms of splenic transcriptomics, while this method was informative of systemic immune remodeling following antibiotic exposure, it ultimately captures the activity of a secondary lymphoid organ and cannot fully recapitulate the local intestinal immune responses at the site of microbial perturbation. Finally, this study did not measure circulating hormone levels, therefore, the observed sex differences in diversity and community composition cannot be directly mechanistically attributed to estrogen signaling and further analyses are needed.

## Conclusion

Overall, this work performed an exploratory study to investigate how continuous antibiotic exposure reshaped the gut microbiome across sexual maturation and how these perturbations influenced downstream host responses in a sex-specific manner. Upon analysis of each omics dataset independently as well as by using an integrative multi-omics framework, the results demonstrated sex-specific correlations at a systems-level across several biological layers including taxonomic and resistome composition, microbial functional pathway potential, host-microbe interactions through cecal metabolomic profiles, and host splenic gene expression assessment. These results provide valuable context to understand systems-level differences observed in males and females following antibiotic exposure and inform sex-specific antibiotic strategies to contribute to more effective antimicrobial stewardship.

## Acknowledgements

This work was supported by resources at the Center for Microbiome Innovation, the Genomics Core at the Sanford Consortium for Regenerative Medicine, the UC San Diego animal care facilities, and the professional veterinary staff for providing their valuable services. This work includes data generated at the UC San Diego IGM Genomics Center utilizing an Illumina NovaSeq X Plus that was purchased with funding from a National Institutes of Health SIG grant (#S10 OD026929).

## Contributions

**Avani Tantry:** Conceptualization; investigation; writing – original draft; writing – review and editing. **Weimiao Long:** Investigation. **Julielam Tran:** Investigation. **Sidhant Rohatgi:** Methodology. **Erika L. Cyphert:** Conceptualization; validation; investigation; writing – original draft; writing – review and editing; supervision; project administration.

## Supplemental Material

Supplemental Figures 1-35 and Supplemental Tables 1-3 are available in a separate document.

## Data Availability

Shotgun Metagenomics Data Accession ID: <u>PRJNA1509482</u>, Untargeted Metabolomics Data Accession ID:, BulkRNASeq Data Accession ID: <u>GSE343025</u>

## Ethics Approval

All experimental procedures were approved by UC San Diego’s Animal Care and Use Committee and performed in strict accordance with NIH Guidelines for the Care and Use of Laboratory Animals.

